# Besides stem canker severity, oilseed rape host genotype matters for the production of *Leptosphaeria maculans* fruiting bodies

**DOI:** 10.1101/2020.08.06.240168

**Authors:** Lydia Bousset, Patrick Vallée, Régine Delourme, Nicolas Parisey, Marcellino Palerme, Melen Leclerc

## Abstract

For fungal cyclic epidemics on annual crops, the pathogen carry-over is an important step in designing disease control strategies. However, it remains particularly difficult to estimate and predict. Plant resistance affects the pathogen development within the epidemics but we lack data on the inter-annual transmission of inoculum. We addressed this question by considering *Leptosphaeria maculans* on 15 oilseed rape genotypes in field during 4 growing seasons. Stem canker severity of host genotypes was visually scored at harvest while the number of fruiting bodies produced on incubated stubble was quantified using an automated image analysis framework. Our results confirm that higher severity at harvest leads to higher fruiting body production and is significantly affected by host genotype and Nitrogen supply. Most interestingly, we show that the production of fruiting bodies is significantly and substantially affected by host genotype, independently of severity at harvest. Tracking individual stems through incubation, we confirm for the first time that the oilseed rape genotype has a direct effect, not only through disease severity. While the genericity of this finding should be investigated on other fungi, this major effect of genotype on inoculum carry-over should be taken into account in models of varietal deployment strategies.

## Introduction

While a key development in terms of disease control has been the breeding and deployment of crop varieties with genetically controlled resistance to pathogens, host qualitative resistance has often not proven durable because most pathogens that pose the greatest threats to crop yields have repeatedly evolved and overcome resistance genes (Burdon *et al*., 2016). Therefore, a critical challenge in plant pathology and epidemiology is to design and implement durable crop protection strategies against rapidly evolving pathogens (Cowger & Brown, 2019). Any mitigation strategy that reduces the size of epidemics and the transfer of inoculum between seasons should also reduce the effective size of pathogen populations, limit the evolutionary potential of pathogens, and increase resistance durability (Bousset & Chèvre, 2013; Zhan *et al*., 2015). Experimental evidence of delayed adaptation has been obtained either by combining quantitative with qualitative resistance (Brun *et al*., 2010: Delourme *et al*., 2014; Lasserre-Zuber *et al*., 2019), by reducing inoculum transmission by the direct action of burying stubble (Daverdin *et al*., 2012), or by removing leaf litter in combination with reduced fungicide application (Didelot *et al*., 2016). The optimal spatio-temporal distribution of host resistance within cultivated landscapes to mitigate disease transmission and prevent the emergence of virulent strains has been addressed through modelling (Lô-Pelzer *et al*., 2010; Papaïx *et al*., 2018; Rimbaud *et al*., 2018; Watkinson-Powell *et al*., 2019) and field experiments (Brun *et al*., 2010; Bousset *et al*., 2018). However, model validation and accurate predictions rely on the availability of data that are difficult and costly to collect at these large scales.

On annual crops, many plant diseases have cyclic epidemics and their dynamics are highly influenced by both temporal and spatial discontinuities, either induced by the climate (e.g. seasonality) or by human actions (e.g. sowing and harvesting) (Zadoks & Schein, 1979; Bousset & Chèvre, 2013; Fig. 1a). The carry-over of inoculum and subsequent dispersal to newly sown fields gives the level of primary inoculum at the beginning of the next cropping season. This carry-over of the pathogen is an important step in designing and predicting the level of success of mitigation strategies, either based on the use of resistant varieties (Marcroft *et al*., 2004a), biocontrol agents (Bailey *et al*., 2004), or preventive management of crop residues (Wherrett *et al*., 2003). However inoculum survival and transmission between seasons remain particularly difficult to monitor in field conditions, and therefore to estimate and predict (Bailey *et al*., 2004; Bousset *et al*., 2015).

**Fig. 1.**
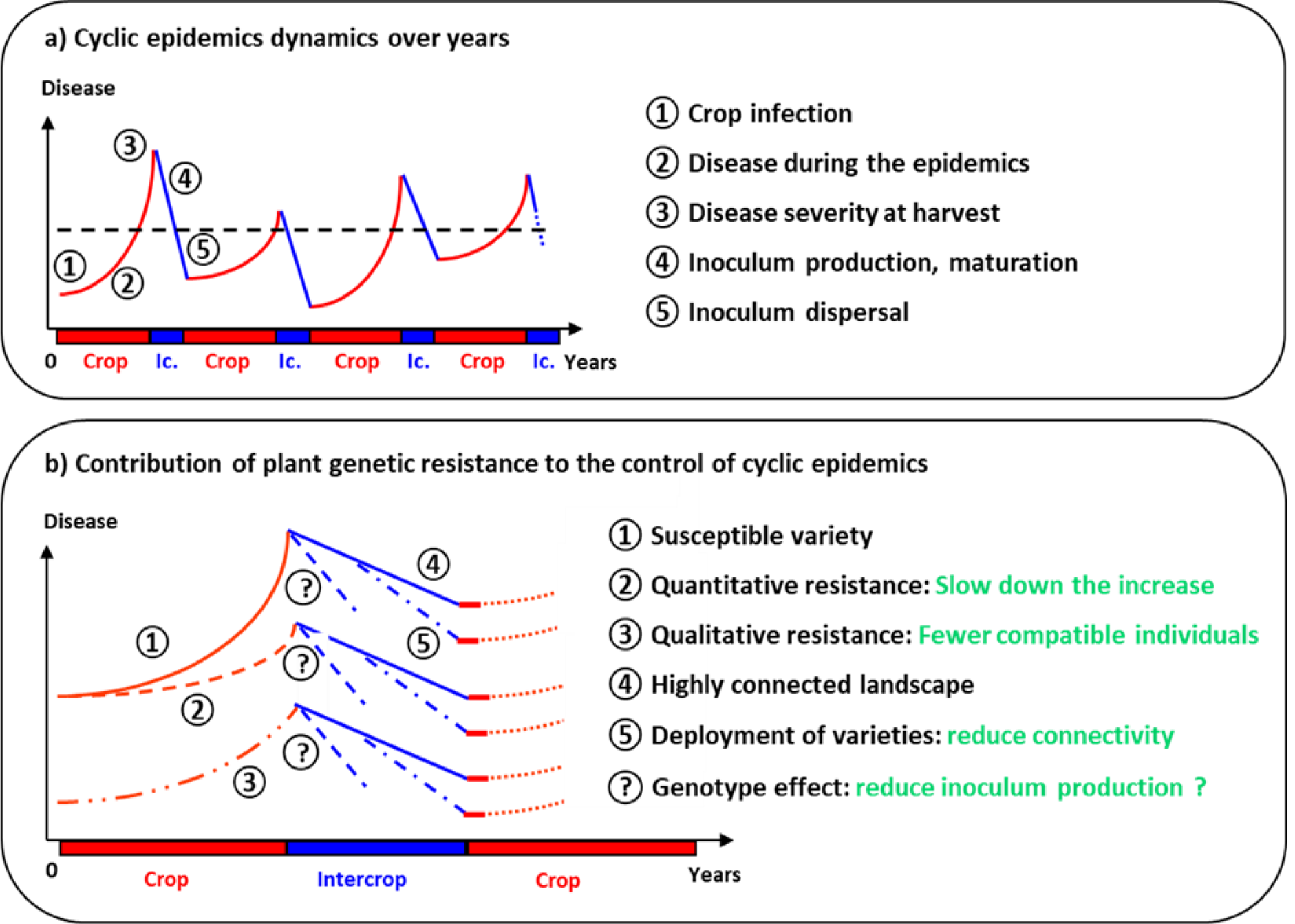
Schematic representation of cyclic epidemics and their control by plant resistance. **a**) Cyclic epidemics are characterized over years by the disease being present in the landscape with a stable long term dynamics (black dashed line), but short term dynamics alternating between epidemic phase (red lines) when the crop is present and decrease (blue lines) during the intercrop (Ic.). **b**) Plant genetic resistance can contribute to the control of cyclic epidemics during the crop with qualitative resistance reducing infection and quantitative resistance reducing the increase. During the intercrop, the selective deployment of genotypes reduces landscape connectivity, and the plant genotype influence the amount of inoculum produced (this study).

Host resistance is usually described by its impact on pathogen life history traits during the epidemic phase, by e.g. infection efficiency, latent period or sporulation (Bruns *et al*., 2012; Delmas *et al*., 2016; Dumartinet *et al*., 2020; Leclerc *et al*., 2019; Bove & Rossi 2020). In short, from an epidemiological point of view, qualitative host resistance reduces the number of compatible infections whereas quantitative host resistance reduces the amplification during the epidemics (Fig. 1b). Following the intercrop, the selective deployment of resistant host genotypes can reduce landscape connectivity (Lô-Pelzer *et al*., 2010; Bousset & Chèvre, 2013; Papaïx *et al*., 2018; Rimbaud *et al*., 2018; Watkinson-Powell *et al*., 2019). Nevertheless, the direct effects of quantitative host resistance on pathogen inter-annual transmission that gives the, generally unknown, initial amount of inoculum at the beginning of the next cropping season remain scarce (Fig. 1b). In *Phytophthora infestans*, a trade-off has been identified between crop infection and survival on tubers meaning that quantitative resistance also affects transmission (Pasco *et al*., 2015). Such trade off was also detected in *Zymoseptoria tritici*, though varietal effects were not investigated (Suffert *et al*., 2018a). In downy powdery mildew, the effect of quantitative resistance was assessed on oospore production, a life history trait related to inoculum inter-annual transmission (Delbac *et al*., 2019).

So far, the most thoroughly studied pathosystem regarding inoculum transmission between seasons is *Leptosphaeria maculans* on oilseed rape. This fungal pathogen initiates epidemics early in the cropping season (in autumn) with stubble-borne ascospores that can spread between fields across the landscape (West *et al*., 2001). It produces phoma leaf spots on host leaves that are observed between autumn and early spring. Then, stem cankers develop from spring to summer, up to the time of harvest, following systemic growth of fungal hyphae from leaf spots to the leaf petiole through xylem vessels, and subsequently to the stem base. The fungus can survive as hyphae in crop stubble, more specifically in stems around crown level, forming two kinds of fruiting bodies: pycnidia and pseudothecia. Pseudothecia can only be formed following sexual reproduction if isolates of opposite mating types co-occur in the same oilseed rape stem. Infected stubble ensures the carry-over of the fungus from one cropping season to the next and serves as the main source of inoculum.

In *L. maculans*, the number of produced pseudothecia, that release initial inoculum ascospores, depends on plant resistance through disease severity at harvest, higher stem canker severities leading to more pseudothecia (McGee & Emmett 1977; Marcroft *et al*., 2004a; Lô-Pelzer *et al*., 2009). Moreover, Marcroft *et al*. (2004b) identified a significant effect of host species and oilseed rape variety on both the visual density of pseudothecia, and the numbers of released ascospores. The authors therefore suggested that a reduced potential for inoculum production could be a trait worth breeding. However, because Marcroft *et al*. (2004b) did not track phoma stem canker severity on individual stems, the observed reduction of fruiting bodies jointly results from reduced canker severity, and potentially from the genotype. Further, by keeping track of individual stem severity and comparing a susceptible cultivar with one having quantitative resistance, Lô-Pelzer *et al*. (2009) were able to see the effect of a genotype with quantitative resistance on the distributions of severities, though they observed no effect of the genotype at a given severity. Afterwards, it has been shown that the production of fruiting bodies on incubated stems can be significantly impacted by the field where they were collected (Bousset *et al*., 2019). However, this study did not allow distinguishing between the effects of genotype and cropping practices. The question of a direct genetic effect on the production of fruiting bodies, and thus temporal transmission of inoculum between seasons, is still unknown for this pathosystem. Addressing this question would require an important amount of work for screening several genotypes with the tedious manual quantification of fruiting bodies. The development of high-precision and high-throughput phenotyping methods and their applications to plant disease epidemiology represent a potential to improve experimental quantification of plant pathogens and improve knowledge and data on pathogen carry-over. For instance, imaging and existing processing algorithms to segment and count regions, offers the prospect to circumvent the long and tedious manual counting (Stewart *et al*., 2016; Karisto *et al*., 2018; Yang & Hong, 2018). In the particular case of *L. maculans*, the automated quantification of fruiting bodies through image processing have already been tackled (Bousset *et al*., 2019), offering a means to overcome the experimental bottleneck and assessing the effect of host quantitative resistance on pathogen carry-over.

In this study, we address the effect of host resistance on the carry-over of inoculum between consecutive years, aiming at disentangling how host genotype might affect fruiting bodies production and the effects of a common cropping practice, i.e. nitrogen fertilization, and disease severity at harvest in the preceding growing season (year). We consider *L. maculans* on oilseed rape as an example pathosystem and use an imaged-based phenotyping method for quantifying fruiting bodies appeared on diseased stems collected in experimental field plots, after an incubation period. We analyse the main drivers of this poorly known life-history trait of the pathogen. We finish by discussing our results and its application to plant disease epidemiology and breeding for resistance.

## Materials and methods

### Experimental fields and contamination of field plots

This experiment was run from 2012 to 2016, with 4 cropping seasons named 1213 to 1516, on the INRAE UE La Motte experimental station located in Le Rheu (48.1°N, 1.5°W), in Brittany, France. Winter *B. napus* genotypes were grown in 5-row plots (2.5 m^2^) and assessed for stem canker (For details of cropping practices see Table S1.1). From 1213 to 1415, the same 15 genotypes were followed (Table 1; S1.2). In 1516, 6 genotypes were added (Table 1). The varieties were chosen to represent a wide range of winter oilseed rape diversity and derived from different breeding programs.

**Table 1.** Numbers of stems for the complete and the three sub datasets (Data1416, Data1216, Data1516) given overall (italicised), but also by Year and Nitrogen fertilization treatment (Nl and Nh) and by Genotype.

| Dataset | Year | Nitrogen | Post process |  |  | Genotype |  |  |  |  |  |  |  |  |  |  |  |  |  |  |  |  |  |  |  |  |
| --- | --- | --- | --- | --- | --- | --- | --- | --- | --- | --- | --- | --- | --- | --- | --- | --- | --- | --- | --- | --- | --- | --- | --- | --- | --- | --- |
|  |  |  | Before | Discard | After | Al | As | Av | Br | Cb | Da | Fa | Go | Gr | Jn | Ko | La | Nh | Po | Yu | Fr | Ja | Mo | Pr | Qu | Ta |
| Complete | All |  | 6758 | 18,3% | 5525 | 390 | 318 | 366 | 325 | 356 | 286 | 338 | 290 | 348 | 247 | 291 | 320 | 363 | 291 | 273 | 114 | 133 | 137 | 76 | 181 | 82 |
|  | 1213 | Nh | 938 | 36 | 902 | 96 | 50 | 70 | 58 | 67 | 64 | 56 | 49 | 54 | 60 | 58 | 44 | 77 | 60 | 39 | - | - | - | - | - | - |
|  | 1314 | Nh | 919 | 48 | 871 | 49 | 59 | 75 | 52 | 88 | 48 | 73 | 42 | 65 | 12 | 31 | 80 | 86 | 70 | 41 | - | - | - | - | - | - |
|  | 1415 | Nh | 826 | 222 | 604 | 72 | 41 | 37 | 15 | 37 | 40 | 57 | 40 | 40 | 43 | 27 | 29 | 57 | 35 | 34 | - | - | - | - | - | - |
|  | 1415 | Nl | 789 | 318 | 471 | 38 | 31 | 30 | 47 | 23 | 25 | 39 | 23 | 32 | 24 | 27 | 36 | 24 | 23 | 49 | - | - | - | - | - | - |
|  | 1516 | Nh | 1626 | 240 | 1386 | 67 | 78 | 77 | 77 | 73 | 57 | 61 | 66 | 71 | 44 | 78 | 71 | 76 | 47 | 59 | 48 | 68 | 91 | 35 | 96 | 46 |
|  | 1516 | Nl | 1660 | 369 | 1291 | 68 | 59 | 77 | 76 | 68 | 52 | 52 | 70 | 86 | 64 | 70 | 60 | 43 | 56 | 51 | 66 | 65 | 46 | 41 | 85 | 36 |
| Data1416 | All |  | 4045 | 25,1% | 3029 | 245 | 209 | 221 | 215 | 201 | 174 | 209 | 199 | 229 | 175 | 202 | 196 | 200 | 161 | 193 | - | - | - | - | - | - |
|  | 1415 | Nh | 826 | 222 | 604 | 72 | 41 | 37 | 15 | 37 | 40 | 57 | 40 | 40 | 43 | 27 | 29 | 57 | 35 | 34 | - | - | - | - | - | - |
|  | 1415 | Nl | 789 | 318 | 471 | 38 | 31 | 30 | 47 | 23 | 25 | 39 | 23 | 32 | 24 | 27 | 36 | 24 | 23 | 49 | - | - | - | - | - | - |
|  | 1516 | Nh | 1208 | 206 | 1002 | 67 | 78 | 77 | 77 | 73 | 57 | 61 | 66 | 71 | 44 | 78 | 71 | 76 | 47 | 59 | - | - | - | - | - | - |
|  | 1516 | Nl | 1222 | 270 | 952 | 68 | 59 | 77 | 76 | 68 | 52 | 52 | 70 | 86 | 64 | 70 | 60 | 43 | 56 | 51 | - | - | - | - | - | - |
| Data1216 | All |  | 3685 | 8,3% | 3379 | 284 | 228 | 259 | 202 | 265 | 209 | 247 | 197 | 230 | 159 | 194 | 224 | 296 | 212 | 173 | - | - | - | - | - | - |
|  | 1213 | Nh | 938 | 36 | 902 | 96 | 50 | 70 | 58 | 67 | 64 | 56 | 49 | 54 | 60 | 58 | 44 | 77 | 60 | 39 | - | - | - | - | - | - |
|  | 1314 | Nh | 919 | 48 | 871 | 49 | 59 | 75 | 52 | 88 | 48 | 73 | 42 | 65 | 12 | 31 | 80 | 86 | 70 | 41 | - | - | - | - | - | - |
|  | 1415 | Nh | 826 | 222 | 604 | 72 | 41 | 37 | 15 | 37 | 40 | 57 | 40 | 40 | 43 | 27 | 29 | 57 | 35 | 34 | - | - | - | - | - | - |
|  | 1516 | Nh | 1002 | 206 | 1002 | 67 | 78 | 77 | 77 | 73 | 57 | 61 | 66 | 71 | 44 | 78 | 71 | 76 | 47 | 59 | - | - | - | - | - | - |
| Data1516 | All |  | 3286 | 18,5% | 2677 | 135 | 137 | 154 | 153 | 141 | 109 | 113 | 136 | 157 | 108 | 148 | 131 | 119 | 103 | 110 | 114 | 133 | 137 | 76 | 181 | 82 |
|  | 1516 | Nh | 1626 | 240 | 1386 | 67 | 78 | 77 | 77 | 73 | 57 | 61 | 66 | 71 | 44 | 78 | 71 | 76 | 47 | 59 | 48 | 68 | 91 | 35 | 96 | 46 |
|  | 1516 | Nl | 1660 | 369 | 1291 | 68 | 59 | 77 | 76 | 68 | 52 | 52 | 70 | 86 | 64 | 70 | 60 | 43 | 56 | 51 | 66 | 65 | 46 | 41 | 85 | 36 |

For the four cropping seasons, Nitrogen supply was high (Nh) according to normal practices in conventional cultivation to achieve yield potential of modern winter oilseed rape varieties (Bouchet et al., 2014) ((Table S1.3). In addition, for 1415 and 1516 seasons, the panel was replicated with normal (Nh) and low (Nl) Nitrogen availability. In order to limit the amount of mineral N in soil in the experimental plots, no organic matter was spread on the fields for 3 years before the trials and the previous crops were grown under a low input management system.

To ensure homogeneous disease infection, contaminated stems collected from the previous season were scattered throughout the plots in each cropping season at a density of two stems per m^2^ when the crop was at the two to three leaf growth stage. To promote fruiting bodies maturation and spore release, sprinklers were run for 2mm irrigation in mornings and afternoons from plant emergence to the apparition of first leaf spots. No fungicides were applied throughout the season. Phoma stem canker severity was assessed 2-3 weeks before crop maturity on a 1 to 6 scale as follows: S1 = no disease, S2a = 1-5%, S2b – 6-25%, S3 = 26-50%, S4 = 51-75%, S5 = 76-99%, S6= 100% of crown cross section with phoma stem canker symptoms. The G2 aggregated disease index was calculated as follows (Aubertot et al. 2004):

G2 = (0 x n_1_+1 x n_2_+3 x n_3_+5 x n_4_+7 x n_5_+9 x n_6_)/N with n_i_ being the number of stems in severity class S_i_, respectively and N the total number of stems.

The climate of the experimental area is oceanic, and meteorological data were obtained from the INRA CLIMATIK database, for Le Rheu weather station, on an hourly basis. Cumulative temperature (Fig. S1ab), rainfall (Fig. S1cd), and number of days that are favourable for pseudothecia maturation were calculated (Fig. S1e). A day was considered favourable if the mean temperature was between 2 and 20°C and if the cumulative rainfall over the previous 11 days before (including the day in question) exceeded 4 mm (Aubertot *et al*., 2006; Lô-Pelzer *et al*., 2009). Given these parameter values, 64 favourable days were required for 50% of pseudothecia to reach maturation.

### Handling of stubble

Following assessment of phoma stem canker severity pieces of stem comprising the crown and upper 10 cm were selected as described in Bousset *et al*. (2019). This sampling was constrained by the availability of diseased stems. A maximum of 30 stems in each of the 6 severity classes was randomly selected from each field while when fewer than 30 stem were available they were all kept. To keep track of phoma canker severity of stem pieces throughout the experiment, two 5 mm diameter holes were drilled, and stem pieces were grouped by 5 on two wooden BBQ sticks painted in blue and labelled with a barcode (Bousset *et al*. 2019).

Over summer, the selected stem pieces were incubated in field conditions at INRAE Le Rheu, on a 1:1:1 mix of sand, peat and compost, with moisture and temperature depending on the local climate and natural rain only. In autumn, these stem pieces were placed in experimental plots of winter oilseed rape and were incubated further. Following maturation, which was indicated by the end of the appearance of new leaf spots on susceptible plants in the field where stem pieces were incubated, the stem pieces were washed, dried and stored (Table 1). All stem pieces from a given cropping season were incubated in the same time and location, starting in the summer following harvest.

### Image acquisition, processing and post-processing

To quantify fruiting bodies appeared on diseased stem pieces, we used the same image-based phenotyping settings as in Bousset *et al*. (2019). In short, a picture of each group of 5 dry stems was taken with the barcoded label, placed on a glass plate, 16 cm above a blue background (PVC sheet Lastolite Colormatt electric blue). Two FotoQuantum LightPro 50 × 70cm softboxes were placed on both sides of the stem pieces with 4 daylight bulbs each (5400K, 30W) within the lower 45° angle. Pictures were taken with a Nikon D5200 with an AF-S DX Micro Nikkor 40mm 1:2.8G lens, on a Kaiser Repro stand, with a wired remote control. Aperture was set at F22 for maximal depth of field, iso 125, daylight white balance. Pictures were saved as RGB images with a resolution of 6000 × 4000 pixels.

Picture pre-processing consisted in reading the barcode to rename the files, splitting each digital image in several new images, each containing only one stem segmented with an unsupervised method. Finally, each new image was cropped to keep only the 5 cm portion on the crown end of the stem, i.e. where pseudothecia are mainly located.

Then, stem pixels were classified either in state state F (fruiting bodies) or in state S (stem) using a supervised machine learning algorithm. As learning and testing data used by Bousset et al. (2019) already included some images collected for this study, we did not re-train the algorithm for processing our images. Afterwards, we conducted a post processing of predictions through a computer-assisted expert curation using a graphical user interface to assign a post-processed state, i.e. “correct” or “incorrect”, to each processed image by visually evaluating the predicted image compared to the original one (Bousset *et al*. 2019).

### Statistical analyses

We analysed the effects of genotype, year and nitrogen fertilization on stem canker severity using a proportional odds model, an ordered logit regression model that handle ordinal data. The design of the experimental data only allowed us to estimate and test the first order effects and the interaction between genotype and nitrogen fertilization.

Then, the density of fruiting bodies detected on images was analysed using the post-processed images validated by the curator (state “correct”). For each image *i* we considered the number of pixels in states F (fruiting bodies) and S (stem), i.e. *n*_*F,i*_ and *n*_*S,i*_, and used a likelihood function based on *n*_*F,i*_ ∼ *B*[(*n*_*F,i*_ + *n*_*S,i*_,), *p*] with a logit link function to build Generalised Linear Models and analyse the effects of selected factors with Wald tests. We first considered the complete unbalanced dataset that only enables the identification and test of first order effects (i.e. year, severity prior incubation, plant genotype and nitrogen fertilization) and the interactions between genotype and nitrogen fertilization, and stem severity with nitrogen fertilization (Table 1). Second, in order to assess more second and third order effects (i.e. interactions) we split the 4 cropping seasons data into 3 datasets : Data1416 (2 years, 2 nitrogen fertilization levels, 15 genotypes), Data1216 (4 years, 15 genotypes, all on high level of fertilization), and Data1516 (21 genotypes and 2 nitrogen fertilization levels all during the 1516 year). Note that for all these subdatasets the severity prior incubation (7 levels) was also included in the regression models.

The whole image processing was developed in Python (Van Rossum & Drake, 1995) whereas statistical analyses were performed using R (R Core Team, 2019).

### Prediction of inoculum potential

The production of inoculum in a population of infected host plants was simulated by coupling two stochastic processes. We first considered 10 experimental plots of 100 plants, whose severity drawn from a Multinomial distribution with event probabilities corresponding to those estimated with the ordered logistic regression model. Second, we assumed that the size of each individual stem is 700 000 pixels, i.e. twice the average observed number of pixels per processed image, and for each stem we drew the number of fruiting bodies pixels using a Binomial distribution, with probability parameter previously estimated in the GLM model. In order to illustrate the importance of host genotype on fruiting bodies production we compared the simulations outputs when considering or not host genotype.

## Results

### Analysis of severity data

The observed distribution of canker severities was assessed at harvest, before the sampling of stems selected for incubation (See Materials and Methods). The proportional odds regression model on the complete ordinal data set pointed out a significant effects (p-values <0.05) of the genotype, the year, the fertilisation and finally the interaction between genotype and fertilisation on canker severity at harvest (Table 2; Fig. 2; Fig. S2.1; Fig. S2.2).

**Table 2.** Analysis of deviance for the observed distribution of canker severities at harvest

| Factor | Df | Chisq | Pr(>Chisq) |
| --- | --- | --- | --- |
| Year | 3 | 7742 | < 2.2e-16 |
| Nitrogen | 1 | 211 | < 2.2e-16 |
| Genotype | 20 | 8828 | < 2.2e-16 |
| Nitrogen:Genotype | 20 | 1971 | < 2.2e-16 |

**Fig. 2.**
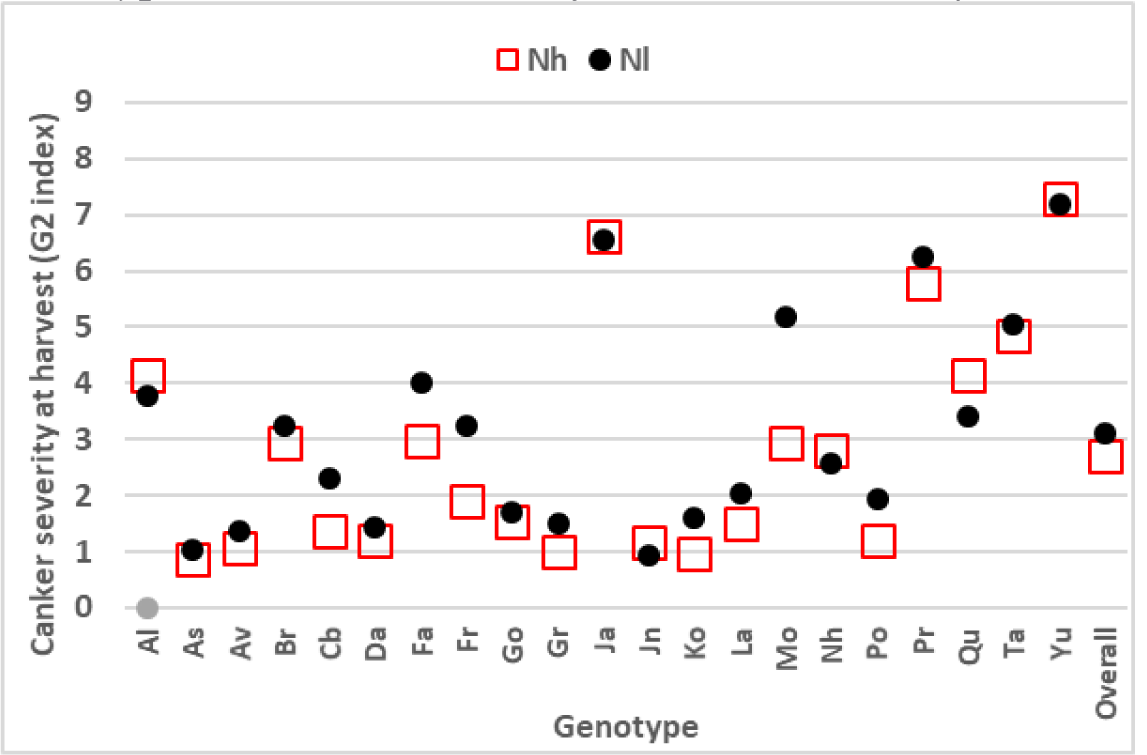
Canker severity (G2 index) on field plots depending on the Nitrogen level and the Genotype. The model was adjusted to the data of 1213 to 1516 Years.

### Year of sampling, genotype and severity had a strong effect on fruiting bodies formation

Whatever the considered dataset, all the tested effects were significant. The year always appeared to be the most influential parameter, explaining about 20% of the explained deviance for all analyses (Table 3). Host genotype was found to explain between 6 and 17% of the explained deviance, indicating a substantial influence of this effect on fruiting bodies formation. Finally, severity of infected stems before incubation explained between 5 and 10 % of the explained deviance. Raw data as well as pairwise comparisons of modalities with least-squares means pointed out differences between genotypes and seasons when stems were collected, 1516 being the one with the least production of fruiting bodies and 1314 the greatest (Fig. 3; Supplementary Information S3).

**Table 3:**
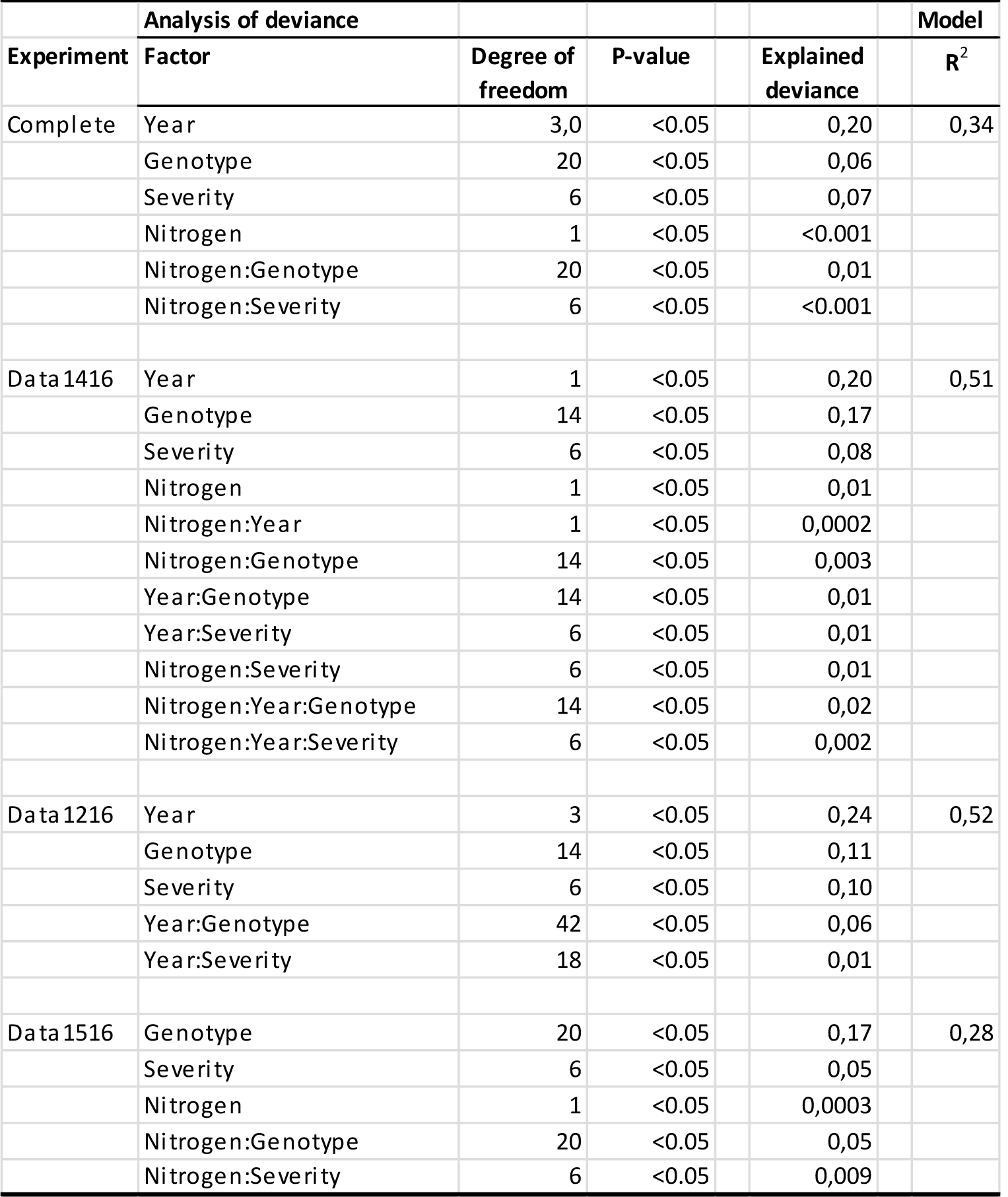
Analysis of deviance for the 3 datasets, with percentage of deviance explained by each factor and by the model

**Fig. 3.**
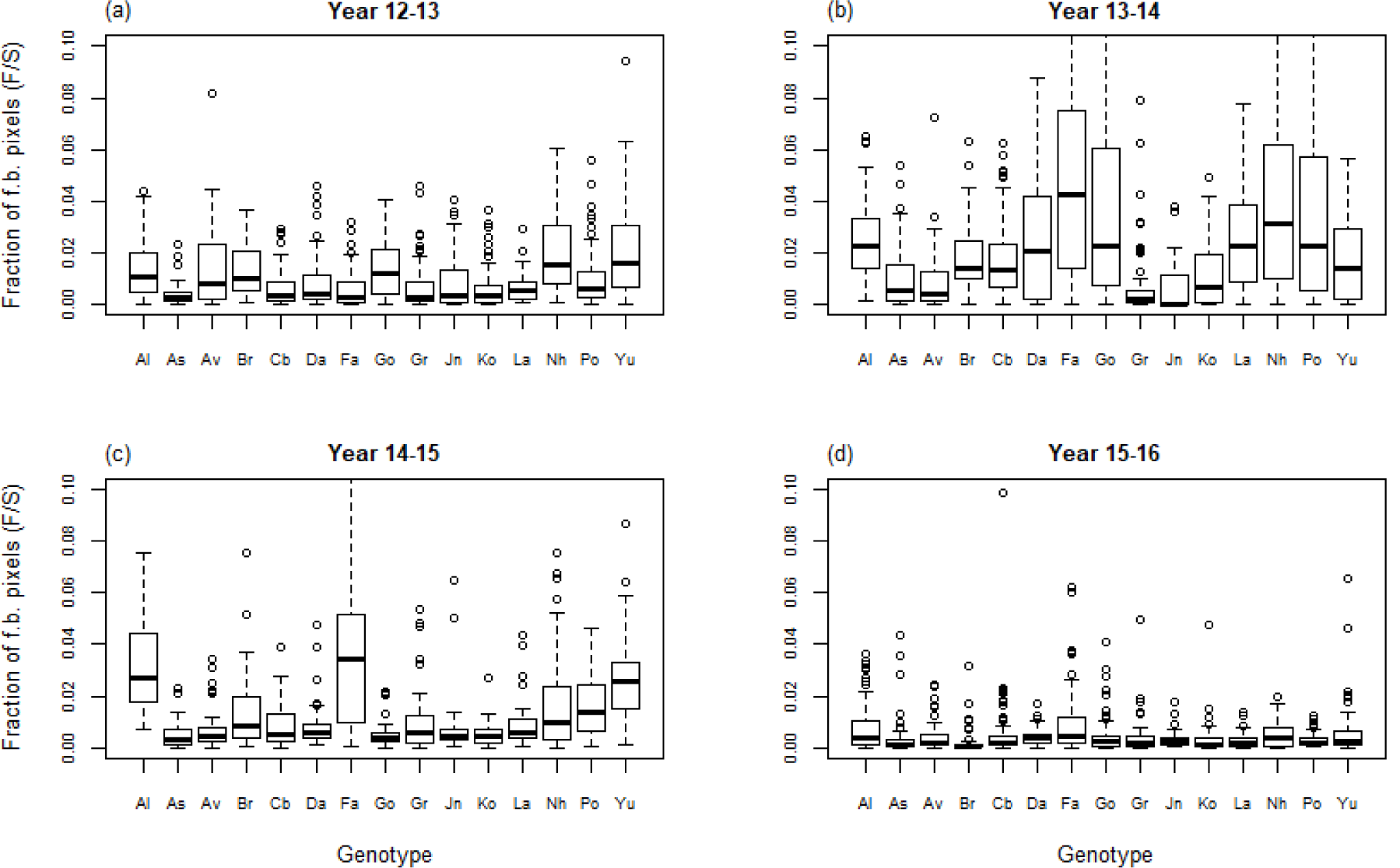
Fraction of fruiting bodies pixels depending on year of sampling (1213, 1314, 1415 and 1516) for the 15 genotypes in the Data1216 dataset.

As already shown by Bousset et al. (2019), the amount of fruiting bodies produced increases with stem canker severity. Stem canker severity classes were ranked from the lowest to highest, except for classes S6 and S5 that were in reversed order (Fig. 4a; Supplementary Information Fig. S3.1; 3.4; 3.7). However, for each severity classes, the fruiting bodies produced depend on the genotype (Fig. 4; 5; Supplementary Information Fig. S4.1; S4.2). Some genotypes like Al and Fa produced greater numbers of fruiting bodies at each severity class than genotypes like As and Av. Noteworthy, here the stem pieces have been individually tracked, so the production of a genotype can be compared at each severity class. One can also note that a severely infected genotype like Yu (having no stem in classes S1 and S2a) is not the one producing the greatest amount of fruiting bodies at a given severity class. On the opposite, a mildly infected genotype like Da (having no stem in classes S5 and S6) produces more fruiting bodies at a given severity class than e.g. genotypes As or Av.

**Fig. 4.**
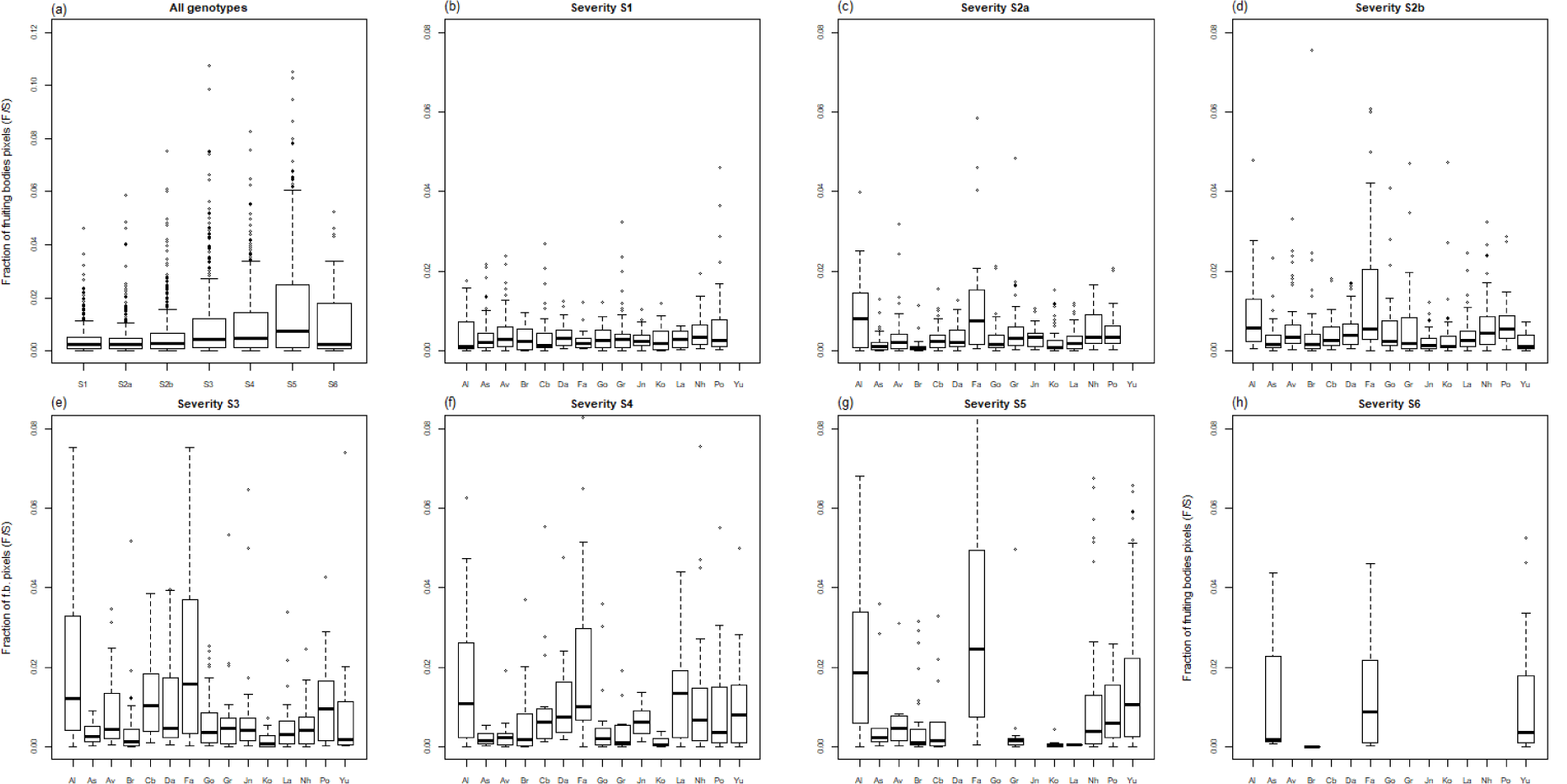
Fraction of fruiting bodies pixels depending for the S1 to S6 severity classes in the Data1416 dataset. a) For all genotypes; b) to h) per genotype for each of the severity classes.

**Fig. 5.**
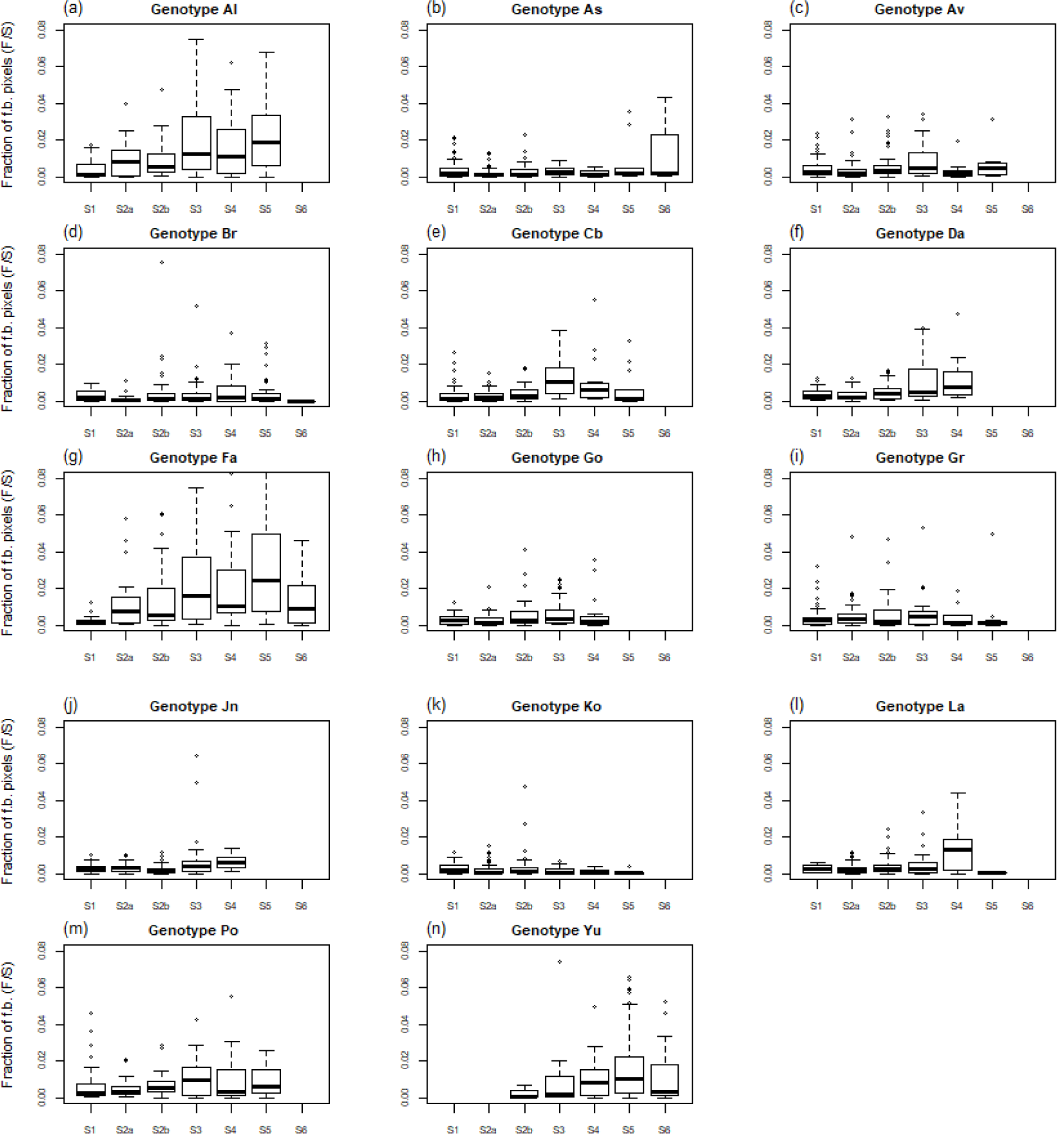
Fraction of fruiting bodies pixels depending on the severity before incubation exemplified for the 15 genotypes in the Data1416 dataset. This result was observed for all datasets (Supplementary Information Fig. S4.1; S4.2).

Because several genotypes did not occur either in the first two (S1-2) or the last two (S5-6) severity classes it was not possible to estimate all the coefficients of the genotype x severity interaction on the all dataset. However, when considering the genotypes for which this interaction was identifiable, we found a significant genotype x severity interaction, suggesting that the amount of produced fruiting bodies by each severity class may change with host genotypes.

### Nitrogen fertilization of the crop has a significant but minor effect on fruiting bodies formation

The effect of nitrogen fertilization significantly affected stem severity (Table 2). Perhaps most interestingly, our analysis also pointed out a significant interaction with host genotype, meaning that the way the level of symptoms changes with Nitrogen supply is not homogeneous across genotypes (Fig. 2).

The analyses of fruiting bodies production exhibited significant effects of fertilization as well as significant interaction with host genotype, host severity and the cultural season. However, as shown by the little deviance explained by these first, second and third order factors, nitrogen fertilization had a minor influence on fruiting bodies production (Table 3; Fig. 6). Nitrogen explained below 1% of the deviance in the Data1516 dataset, and only the interaction Nitrogen:Genotype explained 5% of the deviance (Table 3).

**Fig. 6.**
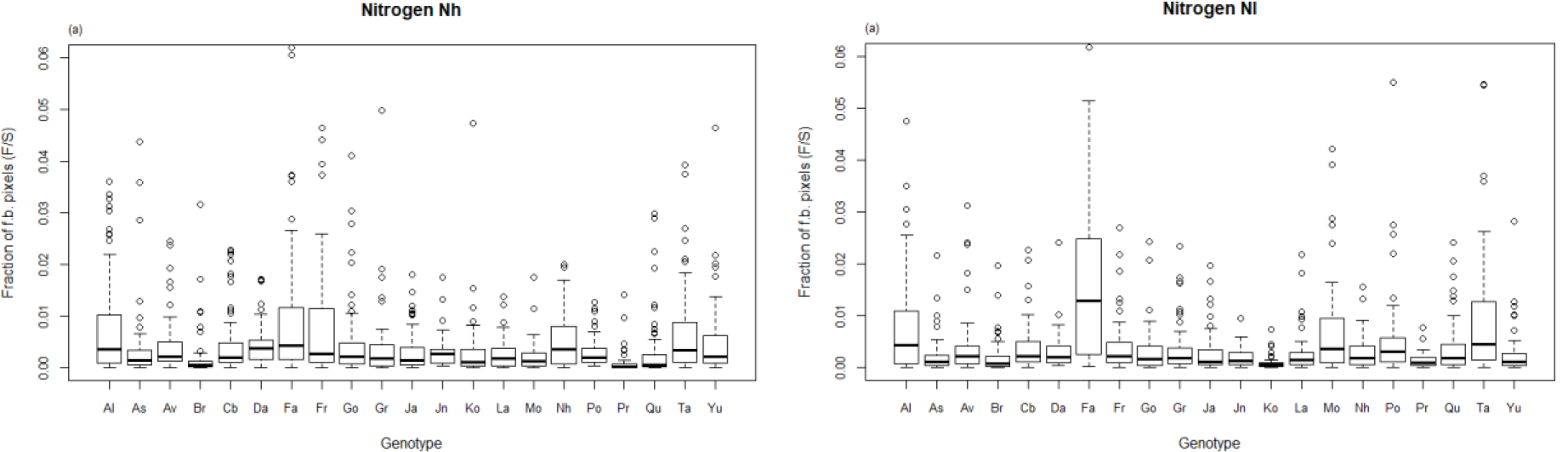
Predicted fraction of fruiting bodies pixels depending on the reduced (Nl) or normal (Nh) Nitrogen fertilization of the crop for the 21 genotypes in the Data1516 dataset.

### Simulated fruiting bodies formation

The stochastic simulation of fruiting bodies production based on fitted event probabilities (proportional odds and generalized linear models) allowed us to predict the potential level of inoculum produced by field plots for each Genotype x Year x Nitrogen fertilization case (Table S5.1; S5.2; S5.3; S5.4). Simulations outputs illustrate the variability of fruiting bodies formation and confirm the importance of host genotype on this life-history trait of the fungal pathogen (Fig. 7; Fig. S5.1).

**Fig. 7.**
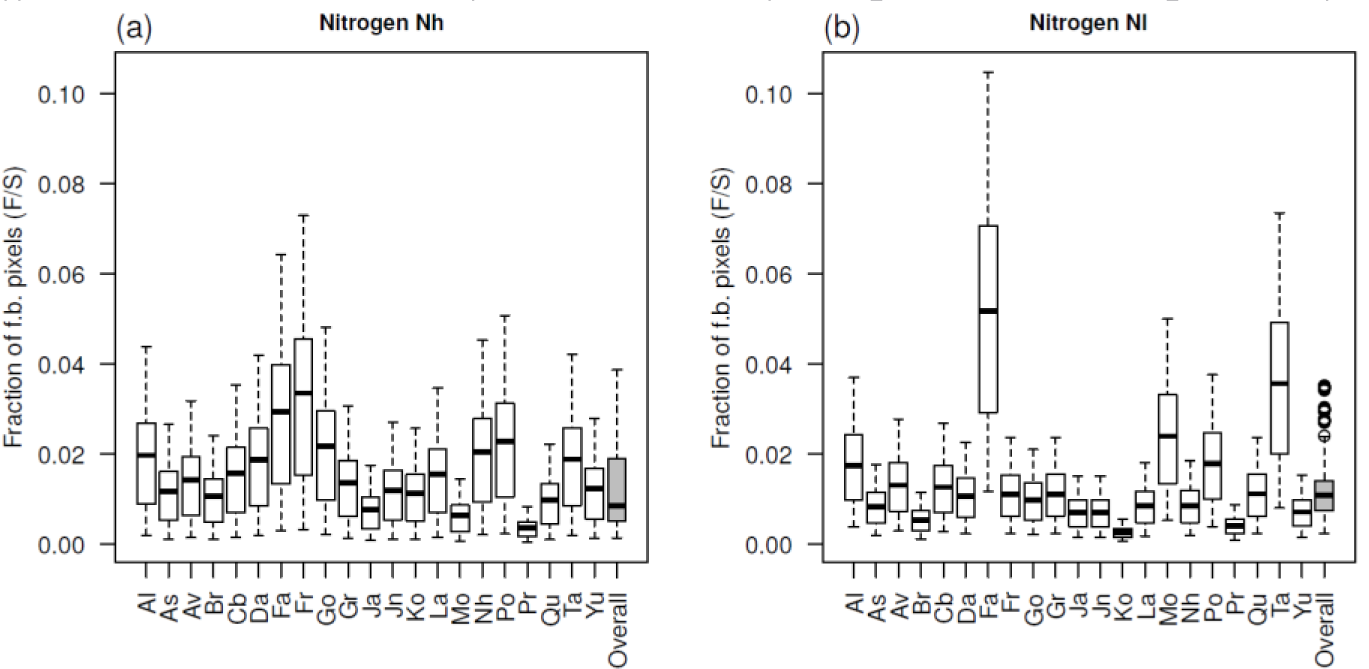
Boxplot of simulated numbers of fruiting bodies pixels depending on the Nitrogen level (Nh and Nl) and on the Genotype (21 varieties). The effect of Genotype was taken into account both for the distribution of stems in canker severity classes and on the prediction of fruiting bodies pixels, adjusting the model on observed data. Red bars are values without the Genotype effect. Data are means of 10 simulated field plots with 100 plants of 700 000 pixels.

When the simulated fruiting bodies production is plotted over canker severity at harvest, one can see that among the genotypes, either susceptible (high canker severity) or with higher levels of quantitative resistance (low canker severity), the pixels of fruiting bodies are variable (Fig. 8). A very susceptible genotype is not always the one leaving the greatest amount of fruiting bodies. Moreover, genotypes with higher levels of quantitative resistance have low canker severity at harvest but are still diverse with regard to fruiting bodies production (Fig. 8).

**Fig. 8.**
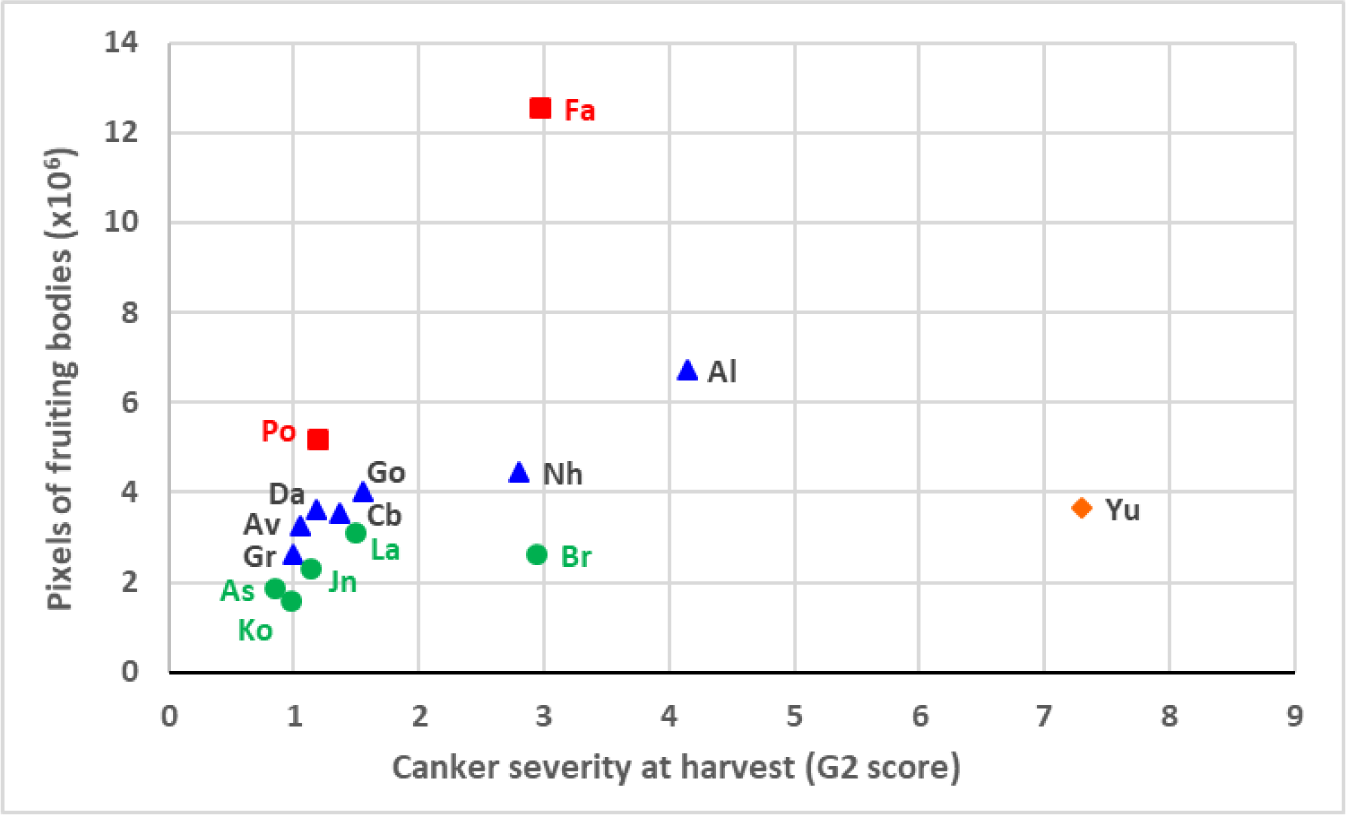
Average simulated numbers of fruiting bodies pixels depending on the canker severity at harvest (G2 index) for the 15 Genotypes over all years. In simulations, the effect of Genotype was taken into account both for the distribution of stems in canker severity classes and on the prediction of fruiting bodies pixels, adjusting the model on observed data. Data are means of 10 simulated field plots with 100 plants of 700 000 pixels. Among the genotypes, either susceptible (high canker severity) or with higher levels of quantitative resistance (low canker severity), the pixels of fruiting bodies are variable. A very susceptible genotype (orange diamond) is not the one leaving the greatest amount of fruiting bodies.

## Discussion

By combining field experiments and an image-based phenotyping method allowing the automated quantification of fruiting bodies on individual stems, we were able to disentangle the role played by the Year, the Genotype, the Severity and finally the Nitrogen fertilization. For the first time, we confirmed that the oilseed rape genotype has a direct effect, not only through disease severity on a seldom-informed trait of the fungal pathogen life cycle. The effect of host genotype on both the visual density of pseudothecia and the numbers of released ascospores had already been observed by Marcroft *et al*. (2004b). However, with their experimental design used, the authors could not clearly distinguish the effect of stem canker severity from the effect of host genotype. Then, Lô-Pelzer *et al*. (2009) followed by Bousset et al. (2019) respectively showed the effect of host quantitative resistance on disease severity and field location on fruiting bodies production. Yet, these studies still not enable to disentangle the role of host genotype, cropping practices as well as the between-season variability. In our study, because all genotypes were planted in the same field and later incubated at the same place, we could exclude confusion with an effect of the environment. Furthermore, as stems were tracked individually, we could separate the effect of severity from the effect of genotype.

The demonstrated variability among host genotypes may be partially explained by differences in host phenology. From the biotrophic and asymptomatic presence of the fungus in the stem, visible cankers appear progressively when crop matures. Moreover, the delay between infection and symptom appearance (i.e. asymptomatic phase) is known to vary between host-plants in a population and the distribution of the incubation period could also change with some variables like the host-age when the infection occurs (Leclerc *et al*., 2014). When collecting disease data from genotypes differing in phenology, the sampling can occur at different stages in these processes, and thus introduce more variability in the relationship between the visual phoma stem canker severity and the resulting fruiting bodies produced. In oilseed rape stems, pathogen load, i.e. the amount of fungal mycelium, remains low throughout winter and starts to increase as *L. maculans* changes from biotrophic to necrotrophic after host flowering (Gervais *et al*., 2017). Nevertheless, in our study neither the time of flowering nor the time between flowering and harvest appeared as a major determinant of the genotype effect (Table S1.2). A more precise quantification of mycelium, by qPCR or by the visualisation of cankers through non-destructive imaging like MRI, could be helpful to better understand the mechanisms involved in the genotype effect.

Genotype effect could be due to a difference in the amount of nutrients available for the fungus. Indeed, in *P. infestans*, the most severely affected plants during the cropping season were the least prone to the intercrop survival of the pathogen on tubers. As quantitative host resistance reduces disease severity during the cropping season, it thus seems to promote intercrop survival (Pasco *et al*., 2015). However, in our study a severely infected genotype like Yu (having no stem in classes S1 and S2a) is neither the one producing the greatest nor the least amount of fruiting bodies at a given severity class. On the opposite, a mildly infected genotype like Da (having no stem in classes S5 and S6) produced more fruiting bodies at a given severity class than e.g. genotypes As or Av. Quantitative host resistance does affect inoculum production by decreasing disease severity, as number of pseudothecia on stems increase with phoma stem canker severity (McGee & Emmett 1977; Marcroft *et al*., 2004a; Lô-Pelzer *et al*., 2009; Bousset *et al*., 2019). In our study, the genotypes Av, Cb, Da, Gr, Jn, Ko had higher than average levels of quantitative resistance on stem canker severity (Table S1.2; Fopa Fomeju *et al*., 2015; Kumar *et al*., 2018), but they did not show any specific trend regarding fruiting bodies production at a given severity class (Fig. 4; 5). Further, our results pointed the significant effect of Nitrogen supply on both disease severity at harvest and fruiting bodies production, though the magnitude of this effect was limited on the latter pathogen life-history trait. It could be worth considering this for breeding cultivar for low-input systems. Genetic analyses are needed to identify the determinants underlying fruiting bodies production in the genotypes. So far, plant resistance is still only characterised during the epidemics, e.g. in grapevine (Bove & Rossi, 2020). However, pathogen life-cycle stages related to intercrop transmission, like the production of oospores in *Plasmopara viticola*, are starting to be evaluated (Delbac *et al*., 2019). Shall heritability be sufficient, this life-history trait of the pathogen could be worth selecting in breeding schemes for pathosystems where host resistance affects pathogen transmission and survival, such as downy mildew on grapevine or stem canker on oilseed rape, at least by discarding the most fruiting bodies-prone genetic backgrounds.

The highlighted effect of host genotype on inoculum carry-over may also be explained by a bias in mating of the fungal pathogen. Because the fungus is heterothallic, mating only occur when the two mating types are present. A small canker with mycelia of both mating types is thus suitable for mating. In the opposite, if a large canker is caused by only one individual, then no pseudothecia could be produced. The occurrence of Allee effects in fungi, i.e. reduced success of mating at low population density, has started to be investigated under controlled conditions for *Zymoseptoria tritici* (Suffert *et al*., 2018b) but should deserve further investigations. In the particular case of *L. maculans*, mycelial growth in the petiole has started to be investigated (Huang *et al*., 2019), but the precise location of the fungus in stem and its consequences on mating remains unknown. Further studies would be interesting and could rely on the use of GFP-transformed strains in controlled conditions or sequencing methods combined with population genetics analyses in natural monitored epidemics.

In agreement with previous findings (Lô-Pelzer *et al*., 2009; Bousset *et al*., 2019), our study also confirmed the important between-year variability that may be related to variable environmental conditions. We observed higher numbers of fruiting bodies in year 1314 and lowest numbers in year 1516 (Fig. 2). These years are the ones with the highest and lowest rainfall during incubation, respectively (Fig. S1d). While the effects of climatic variables on maturation of the fruiting bodies have been modelled (Aubertot *et al*., 2006), their influences on the pathogen during the cropping season and during the intercrop still deserve further investigations. In particular, besides influencing the development of the host plant, some main climatic variables such as temperature may drive within-host pathogen growth, influence pathogen load at stem base, thus influence mating of the fungus and therefore fruiting bodies production. Linking environmental conditions with host and pathogen development is a current challenge for most pathosystems to improve predictions of epidemics and yields. In the particular case of oilseed rape stem canker, further studies are needed to address the influence of crop growth in real situations as well as intercrop practices of the production of fruiting bodies (McCredden *et al*., 2017).

As plant genotype appeared to be an important driver of the production of inoculum, this should be taken into account in models used to compare strategies for the deployment of varieties in the landscape (Lô-Pelzer *et al*., 2010; Papaïx *et al*., 2018; Rimbaud *et al*., 2018; Watkinson-Powell *et al*., 2019). So far, the effect of qualitative host resistance on infection and some effects of quantitative host resistance on pathogen development are considered. Our results suggest that it would be worth considering the effect of host quantitative resistance on the pathogen during the intercrop. One striking result of our stochastic simulations of fruiting bodies production is that the ranking of the genotypes (Fig. 7) appeared to be very different from the ranking obtained when considering the disease severity at harvest (Fig. 2). For example, the genotypes Ja, Pr and Yu with a high disease severity at harvest (Fig.2) produce fewer fruiting bodies that others (Fig.7). On the opposite, the genotypes Fa, Fr and Po have a moderate disease severity at harvest (Fig.2) but later produce higher amounts of fruiting bodies that others (Fig.7). This finding is likely to change our vision of what is a good resistant cultivar for a durable control of pathogen while mitigating the effects of disease on yield yearly. Such ideotype could be identified using plurennial epidemiological framework that integrate all the components of resistance, in models implemented with data acquired from efficient sensors. This would be helpful to quantify the benefits of breeding less prone to inoculum production and guide further breeding schemes.

## Acknowledgements

We thank UE La Motte for running the cultivation of the experiments. We thank Yannick Lucas, Claude Domin and casual workers for technical assistance. We thank BraCySol biological resource centre (INRA Ploudaniel, France) for providing the seeds used in this study. We are grateful to the INRA CLIMATIK database for the weather data. This work benefited from the financial support of INRA - the French National Institute for Agronomical Research, and from ANR - the French National Research Agency – program AGROBIOSPHERE grant ANR-11-AGRO-003-01 and from the French Association for the Promotion of Oilseed Crops Breeding (PROMOSOL) through the project MOREAZ (Modulation de la réponse de résistance du colza à des agents pathogènes (hernie et phoma) en situation de contrainte azotée).

## Authors’ contributions

LB, PV and RD carried out experiments, ML carried out statistical analyses, LB, ML, NP and MP analyzed pictures. LB conceived and designed the study and prepared the manuscript, read and approved by all authors. The authors declare the absence of conflict of interest.

## Supplementary information

**S-1 Assessment of climatic conditions during crop growth and pseudothecia maturation, details about cropping practices and about the genotypes used**

**Fig. S1** Assessment of climatic conditions during crop growth and pseudothecia maturation in four cropping seasons 2012-2013 (Inc. of 1213) to 2015-2016 (Inc. of 1516). **a**). Mean daily temperature was cumulated over the course of each experiment from sowing to harvest. **b**). Mean daily temperature was cumulated over the course of each experiment from start to end of stem incubation. **c**). Cumulative rainfall during crop growth (from sowing to harvest). **d**). Cumulative rainfall from start to end of stem incubation E. Cumulative numbers of days favourable for the maturation of pseudothecia. A day was considered favourable if the mean temperature was between 2 and 20C and if the cumulative rainfall over the previous 11 days beforehand (including the day in question) exceeded 4 mm (Aubertot et al. 2006; L-Pelzer et al. 2009). Given these parameter values, 64 favourable days are required for 50% of pseudothecia to reach maturation. Meteorological data were obtained from the INRA CLIMATIK database, for Le Rheu weather station, on an hourly basis.

**Table S1.1** Dates of cropping practices and stem incubation for the three datasets (Data1416, Data1216, Data1516). Sums of degree days were calculated for the whole season, or for the autumn (sowing to 31 December), pre flowering (1^st^ January to flowering) and post flowering (flowering to harvest) periods

**Table S1.2:** Oilseed rape varieties with their breeding origin, seed quality, flowering dates and disease severity estimated by Kumar *et al*. 2018.

**Table S1.3** Nitrogen fertilisation timing and amounts (Units / Hectare) in the years (1213 to 1516) of the experiment. In 1415 and 1516 the experiment was replicated with high (Nh) and reduced (Nl) Nitrogen.

**S-2 Canker severity at harvest**

**Fig. S2.1** Distributions of stems in the 7 canker severity classes (S1 to S6, shades of grey) at harvest depending on the year (1213 to 1516) and on the Genotype (21 varieties).

**Fig. S2.2** Mean canker severity at harvest depending on the Nitrogen level (Nh = normal, Nl = reduced) and on the Genotype (21 varieties). Results are least square means following analysis of variance.

**S-3. Summary figures for each of the three datasets and pairwise comparisons of least square means across Severity classes and across Genotypes**

**Fig. S3.1** Graphical examination of the fraction (F/S) of fruiting bodies pixels in states F (fruiting bodies) and S (stem) on 3029 oilseed rape stems after the processing and post-processing of RGB digital images in the Data1416 dataset. a) Histogram of class frequency depending on fruiting bodies fraction ; b-d), Boxplots showing how the fraction F/S changed b) between 2 Years; c) between 7 classes of phoma stem canker Severity before incubation; d) between 15 Genotypes. Phoma stem canker severity prior incubation classes S1 to S6 correspond to increasing proportion of cankered cross section at crown level. Year, disease Severity and Genotype all have a significant effect.

**Table S3.1** The pairwise comparisons of least-square means pointed out a significant difference between the Severity classes in the Data1416 dataset. Results are averaged over the levels of severity for each of the 2 Years and the 2 Nitrogen fertilization levels; Confidence level used: 0.95; P value adjustment: Tukey method for comparing a family of 7 estimates; Significance level used: alpha = 0.05.

**Fig. S3.2** Predicted probability of fruiting bodies pixels by Severity class for each of the 2 Years and the 2 Nitrogen fertilization levels in the Data1416 dataset.

**Table S3.2** The pairwise comparisons of least-square means pointed out a significant difference between the genotypes in the Data1416 dataset. Results are averaged over the levels of Genotypes for each of the 2 Years and the 2 Nitrogen fertilization levels; Confidence level used: 0.95; P value adjustment: Tukey method for comparing a family of 15 estimates; Significance level used: alpha = 0.05.

**Fig. S3.3** Predicted probability of fruiting bodies pixels by Genotype for each of the 2 Years and the 2 Nitrogen fertilization levels in the Data1416 dataset.

**Fig. S3.4** Graphical examination of the fraction (F/S) of fruiting bodies pixels in states F (fruiting bodies) and S (stem) on 3379 oilseed rape stems after the processing and post-processing of RGB digital images in the Data1216 dataset. a) Histogram of class frequency depending on fruiting bodies fraction ; b-d), Boxplots showing how the fraction F/S changed b) between 4 Years; c) between 7 classes of phoma stem canker Severity before incubation; d) between 15 Genotypes. Phoma stem canker severity prior incubation classes S1 to S6 correspond to increasing proportion of cankered cross section at crown level. Year, disease Severity and Genotype all have a significant effect.

**Table S3.3** The pairwise comparisons of least-square means pointed out a significant difference between the Severity classes in the Data1216 dataset. Results are averaged over the levels of severity for each of the 4 Years and the Nh Nitrogen fertilization level; Confidence level used: 0.95; P value adjustment: Tukey method for comparing a family of 7 estimates; Significance level used: alpha = 0.05.

**Fig. S3.5** Predicted probability of fruiting bodies pixels by Severity class for each of the 4 Years and the Nh Nitrogen fertilization level in the Data1216 dataset.

**Table S3.4** The pairwise comparisons of least-square means pointed out a significant difference between the genotypes in the Data1216 dataset. Results are averaged over the levels of Genotypes for each of the 4 Years and the Nh Nitrogen fertilization level; Confidence level used: 0.95; P value adjustment: Tukey method for comparing a family of 15 estimates; Significance level used: alpha = 0.05.

**Fig. S3.6** Predicted probability of fruiting bodies pixels by Genotype for each of the 4 Years and the Nh Nitrogen fertilization level in the Data1416 dataset.

**Figure S3.7** Graphical examination of the fraction (F/S) of fruiting bodies pixels in states F (fruiting bodies) and S (stem) on 2677 oilseed rape stems after the processing and post-processing of RGB digital images in the Data1516 dataset. a) Histogram of class frequency depending on fruiting bodies fraction ; b-d), Boxplots showing how the fraction F/S changed b) between 4 Years; c) between 7 classes of phoma stem canker Severity before incubation; d) between 21 Genotypes. Phoma stem canker severity prior incubation classes S1 to S6 correspond to increasing proportion of cankered cross section at crown level. Disease Severity, and Genotype all have a significant effect.

**Table S3.5** The pairwise comparisons of least-square means pointed out a significant difference between the Severity classes in the Data1516 dataset. Results are averaged over the levels of severity for the 1516 Year and the 2 Nitrogen fertilization level; Confidence level used: 0.95; P value adjustment: Tukey method for comparing a family of 7 estimates; Significance level used: alpha = 0.05.

**Figure S3.8** Predicted probability of fruiting bodies pixels by Severity class for the 1516 Year and the 2 Nitrogen fertilization level in the Data1516 dataset.

**Table S3.6** The pairwise comparisons of least-square means pointed out a significant difference between the genotypes in the Data1516 dataset. Results are averaged over the levels of Genotypes for the 1516 Year and the 2 Nitrogen fertilization level; Confidence level used: 0.95; P value adjustment: Tukey method for comparing a family of 21 estimates; Significance level used: alpha = 0.05.

**Figure S3.9** Predicted probability of fruiting bodies pixels by Genotype for the 1516 Year and the 2 Nitrogen fertilization level in the Data1516 dataset.

**S-4. Fraction of fruiting bodies pixels depending on the severity before incubation for each of the genotypes in each of the datasets**.

**Fig. S4.1**. Fraction of fruiting bodies pixels depending on the severity before incubation for the 15 genotypes in the Data1216 dataset.

**Fig. S4.2**. Fraction of fruiting bodies pixels depending on the severity before incubation for the 15 genotypes in the Data1516 dataset.

**S-5 Simulations**

**Table S5.1** Parameters estimates of the proportional odds model with genotype effect used to predict event probabilities in the multinomial draw of simulations

**Table S5.2** Parameters estimates of the proportional odds model without genotype effect used to predict event probabilities in the multinomial draw of simulations

**Table S5.3** Parameters estimates of GLM model with genotype effect used to predict the probability in the Binomial draw of simulations

**Table S5.4** Parameters estimates of GLM model without genotype effect used to predict the probability in the Binomial draw of simulations

**Fig. S5.1** Boxplot of simulated numbers of fruiting bodies pixels depending on the Year (2013 to 2016), on the Nitrogen level (Nh and Nl) and on the Genotype (21 varieties). The effect of Genotype was taken into account both for the distribution of stems in canker severity classes and on the prediction of fruiting bodies pixels, adjusting the model on observed data. Red bars are values without the Genotype effect. Data are means of 10 simulated field plots with 100 plants of 700 000 pixels.

