## Supplementary Information for "Besides stem canker severity, oilseed rape host genotype matters for the production of *Leptosphaeria maculans* fruiting bodies"

### S-1 Assessment of climatic conditions during crop growth and pseudothecia maturation, details about cropping practices and about the genotypes used

**Fig. S1** Assessment of climatic conditions during crop growth and pseudothecia maturation in four cropping seasons 2012-2013 (Inc. of 1213) to 2015-2016 (Inc. of 1516). **a**). Mean daily temperature was cumulated over the course of each experiment from sowing to harvest. **b**). Mean daily temperature was cumulated over the course of each experiment from start to end of stem incubation. **c**). Cumulative rainfall during crop growth (from sowing to harvest). **d**). Cumulative rainfall from start to end of stem incubation E. Cumulative numbers of days favourable for the maturation of pseudothecia. A day was considered favourable if the mean temperature was between 2 and 20C and if the cumulative rainfall over the previous 11 days beforehand (including the day in question) exceeded 4 mm (Aubertot et al. 2006; L-Pelzer et al. 2009). Given these parameter values, 64 favourable days are required for 50% of pseudothecia to reach maturation. Meteorological data were obtained from the INRA CLIMATIK database, for Le Rheu weather station, on an hourly basis.

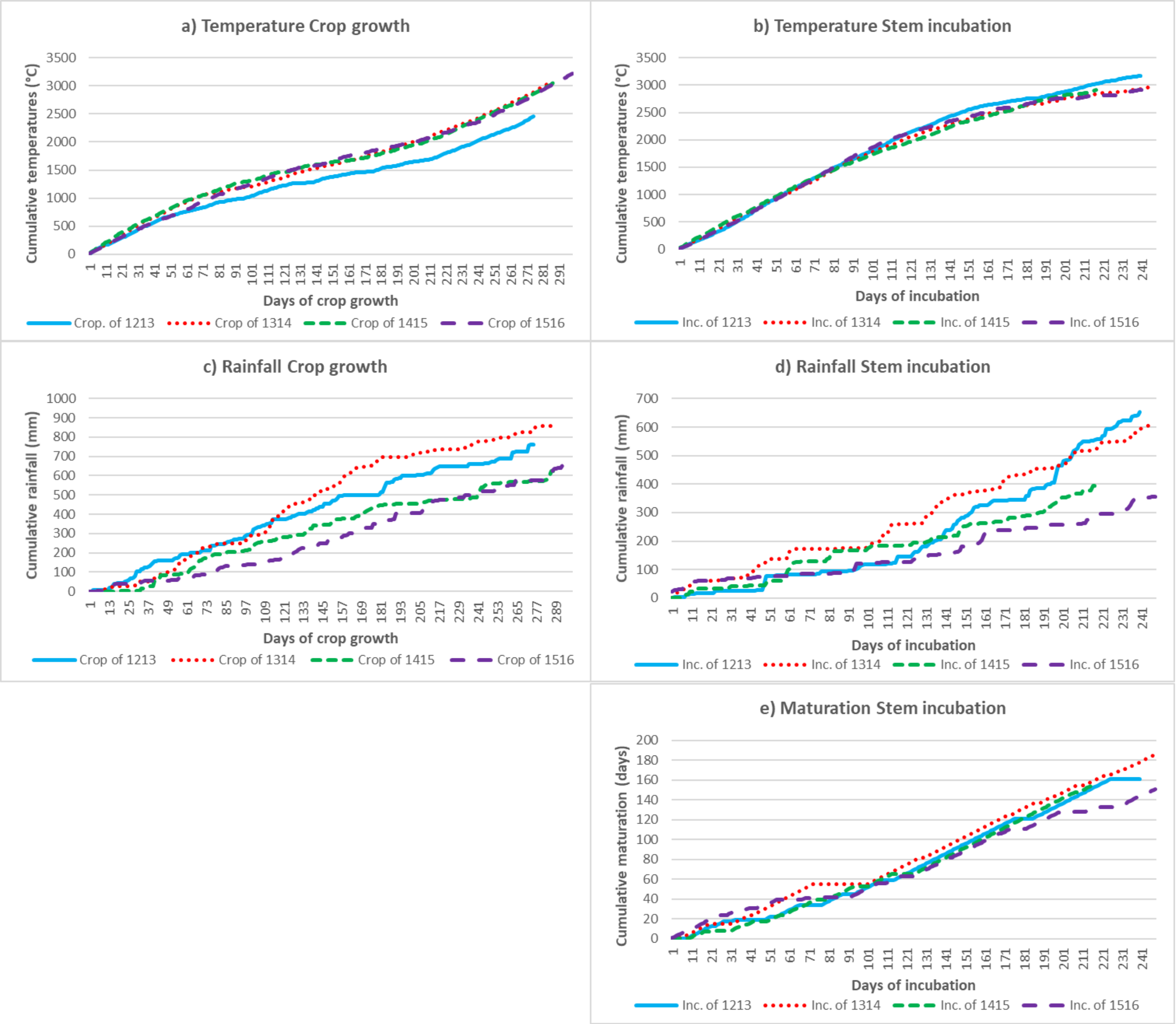

**Table S1.1** Dates of cropping practices and stem incubation for the three datasets (Data1416, Data1216, Data1516). Sums of degree days were calculated for the whole season, or for the autumn (sowing to 31 December), pre flowering (1st January to flowering) and post flowering (flowering to harvest) periods

| Cropping practices | | | Cumulative Temperature (Degre Days) | | | | Cumulative Rainfall (mm) | | | | Stem incubation | | | | |
|---|---|---|---|---|---|---|---|---|---|---|---|---|---|---|---|
| Sowing | Flowering | Harvest | Season | Autumn | Pre. Flo. | Post. Flo. | Season | Autumn | Pre. Flo. | Post. Flo. | Start | End | Deg. days | Rain (mm) | Matur. |
| 10.09.2012 | 08.04/26.04 | 06.06.2013 | 2452 | 1179 | 517 | 549 | 760 | 373 | 245 | 111 | 10.06.2013 | 04.02.2014 | 3174 | 652 | 161 |
| 03.09.2013 | 24.03/11.04 | 16.06.2014 | 3097 | 1371 | 647 | 881 | 857 | 422 | 296 | 121 | 26.06.2014 | 27.02.2015 | 2980 | 616 | 185 |
| 02.09.2014 | 31.03/17.04 | 15.06.2015 | 3048 | 1460 | 575 | 810 | 625 | 282 | 188 | 147 | 26.06.2015 | 28.01.2016 | 2907 | 394 | 156 |
| 02.09.2015 | 22.03/25.04 | 28.06.2016 | 3226 | 1463 | 541 | 900 | 666 | 173 | 232 | 166 | 14.06.2016 | 16.02.2017 | 2962 | 356 | 151 |

**Table S1.2:** Oilseed rape varieties with their breeding origin, seed quality, flowering dates and disease severity estimated by Kumar *et al*. 2018.

| Genotype | Variety | Breeder[a] | Seed quality[b] | Disease[c] | Flowering[d] | | | | | |
|---|---|---|---|---|---|---|---|---|---|---|
| | | | | | 1213_Nh | 1314_Nh | 1415_Nh | 1415_Nl | 1516_Nh | 1516_Nl |
| Al | Alsace 1360 | INRA | ++ | 1,5 | 19-Apr | 1-Apr | 12-Apr | 12-Apr | 12-Apr | 12-Apr |
| As | ES Astrid | Euralis | 00 | 1,0 | 13-Apr | 26-Mar | 7-Apr | 5-Apr | 8-Apr | 6-Apr |
| Av | Aviso | Danisco Seeds | 00 | -1,2 | 16-Apr | 31-Mar | 10-Apr | 7-Apr | 5-Apr | 4-Apr |
| Br | Bristol | Monsanto SAS | 00 | 0,7 | 9-Apr | 26-Mar | 5-Apr | 6-Apr | 8-Apr | 5-Apr |
| Cb | Canberra | Monsanto SAS | 00 | -1,3 | 12-Apr | 24-Mar | 5-Apr | 3-Apr | 1-Apr | 1-Apr |
| Da | Darmor | INRA/Serasem | 00 | -1,1 | 23-Apr | 7-Apr | 15-Apr | 14-Apr | 21-Apr | 20-Apr |
| Fa | Falcon | NPZ | 00 | 0,2 | 13-Apr | 31-Mar | 5-Apr | 3-Apr | 31-Mar | 1-Apr |
| Fr | Frederic | RAPS Gbr | 00 | -0,9 | 12-Apr | 24-Mar | 4-Apr | 3-Apr | 4-Apr | 2-Apr |
| Go | Goeland | Dippe Saatzucht | 00 | -0,7 | 19-Apr | 3-Apr | 13-Apr | 13-Apr | 17-Apr | 11-Apr |
| Gr | Grizzly | RAGT 2n | 00 | -1,5 | 19-Apr | 3-Apr | 13-Apr | 13-Apr | 15-Apr | 13-Apr |
| Jn | Jet-Neuf | Serasem | 0+ | -1,3 | 20-Apr | 3-Apr | 13-Apr | 12-Apr | 15-Apr | 15-Apr |
| Ko | Kosto | Momont | 00 | -1,4 | 22-Apr | 2-Apr | 13-Apr | 13-Apr | 15-Apr | 13-Apr |
| La | Labrador | Momont | 00 | -0,5 | 19-Apr | 3-Apr | 11-Apr | 13-Apr | 14-Apr | 12-Apr |
| Mo | Mohican | CPB | 00 | 0,2 | 16-Apr | 31-Mar | 9-Apr | 9-Apr | 2-Apr | 2-Apr |
| Nh | Nain de Hambourg | INRA | ++ | 0,0 | 17-Apr | 31-Mar | 10-Apr | 10-Apr | 8-Apr | 8-Apr |
| Po | Pollen | Momont | 00 | -0,7 | 15-Apr | 26-Mar | 8-Apr | 9-Apr | 4-Apr | 2-Apr |
| Qu | Quinta | NPZ | 0+ | 2,2 | 24-Apr | 3-Apr | 13-Apr | 13-Apr | 25-Apr | 25-Apr |
| Ja | Jantar | IHAR | 00 | 6,7/8,4[e] | 23-Apr | 2-Apr | 17-Apr | 14-Apr | 20-Apr | 15-Apr |
| Pr | Primor | INRA | 0+ | 5,6/6,8[e] | 17-Apr | 31-Mar | 10-Apr | 9-Apr | 12-Apr | 8-Apr |
| Ta | Tapidor | France | 00 | Susceptible | - | - | 3-Apr | 3-Apr | 1-Apr | 29-Mar |
| Yu | Yudal | Spring korean line | ++ | Susceptible | 8-Apr | 24-Mar | 1-Apr | 31-Mar | 22-Mar | 22-Mar |
| Overall mean | | | | | 16-Apr | 30-Mar | 9-Apr | 8-Apr | 9-Apr | 8-Apr |

[a]Abbreviations are CPB: Cambridge Plant Breeders; IHAR: Instytut Hodowli i Aklimatyzacji Roslin, Poznan; INRA: Institut National de la Recherche Agronomique; NPZ: Norddeutsche Pflanzenzucht Hans-Georg Lembke KG.

[b]Seed quality is ++: Oilseed rape with high erucic acid and high glucosinolate content in the seeds; 0+: Oilseed rape with low erucic acid and high glucosinolate.

[c] Disease is BLUP (Best Linear Unbiased prediction) phoma stem canker disease index estimated by Kumar *et al*. 2018.

[d] Flowering is the day when 50 % of the plants showed 10 % of open flowers on the primary inflorescence.

[c]G2 Disease index (on a 0-9 scale) in 2006 / 2013, from Fopa-Fomeju *et al*. 2015.

**Table S1.3** Nitrogen fertilisation timing and amounts (Units / Hectare) in the years (1213 to 1516) of the experiment. In 1415 and 1516 the experiment was replicated with high (Nh) and reduced (Nl) Nitrogen.

| Year | | Fall[a] | | Spring[a] | |
|---|---|---|---|---|---|
| | | Amount (U/Ha) | Date | Amount | Date |
| 1213 | Nh | 35 U/Ha | 22.08.2012 | 40 U/Ha | 25.02.2013 |
| 1314 | Nh | 30 U/Ha | 20.08.2013 | 40+40 U/Ha | 27.02.2014 / 13.03.2014 |
| 1415 | Nh | none | | 40+40 U/Ha | 02.03.2015 / 20.03.2015 |
| 1415 | Nl | none | | 40 U/Ha | 02.03.2015 |
| 1516 | Nh | none | | 80 U/Ha | 29.02.2016 |
| 1516 | Nl | none | | 60 U/Ha | 29.02.2016 |

[a]Fall fertilisation was manure and spring applications were a liquid fertilizer with a 39 % N solution (50 % urea, 25 % nitrate and 25 % ammonium)

### S-2 Canker severity at harvest

**Fig. S2.1** Distributions of stems in the 7 canker severity classes (S1 to S6, shades of grey) at harvest depending on the year (1213 to 1516) and on the Genotype (21 varieties).

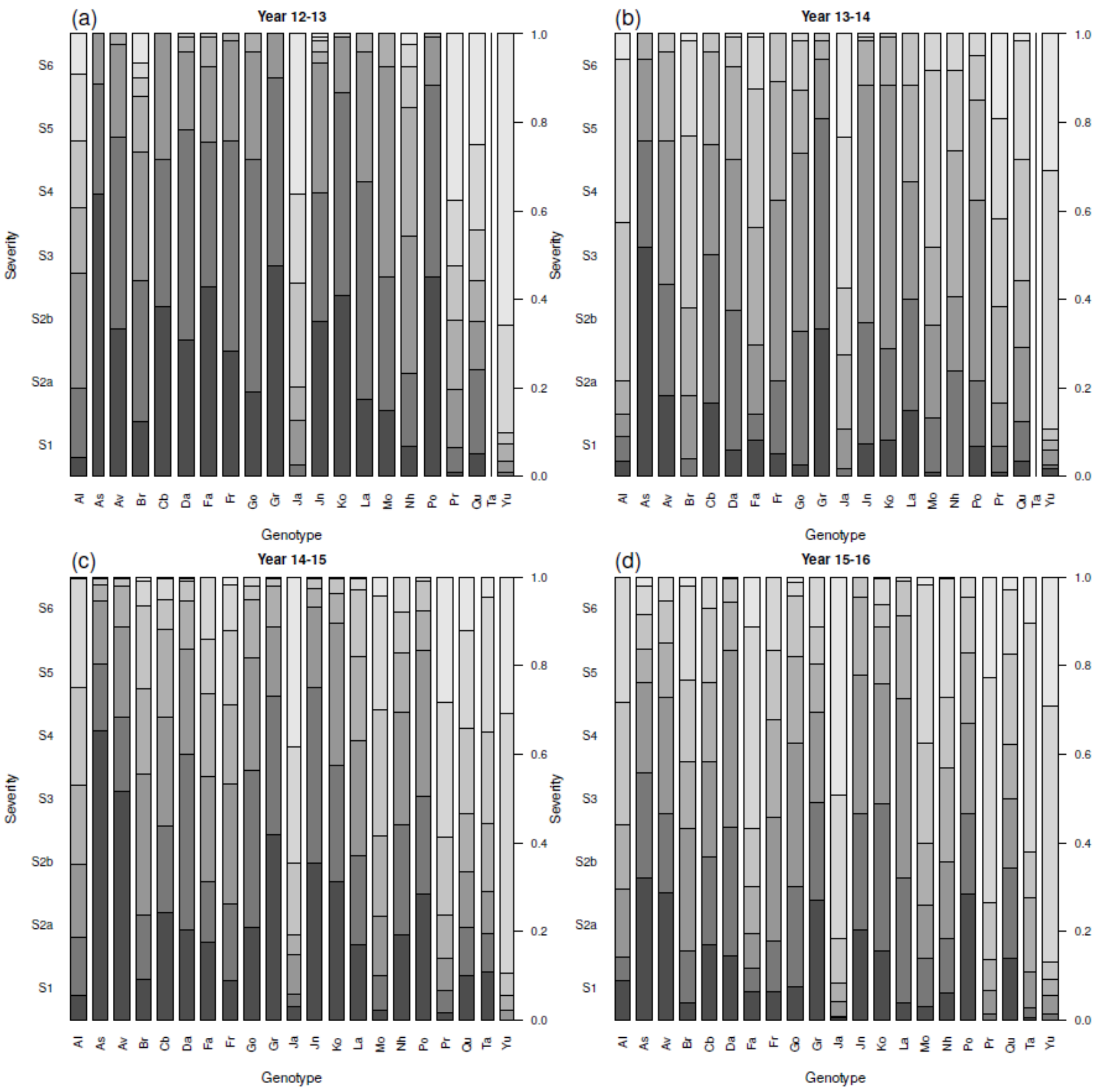

**Fig. S2.2** Mean canker severity at harvest depending on the Nitrogen level (Nh = normal, Nl = reduced) and on the Genotype (21 varieties). Results are least square means following analysis of variance.

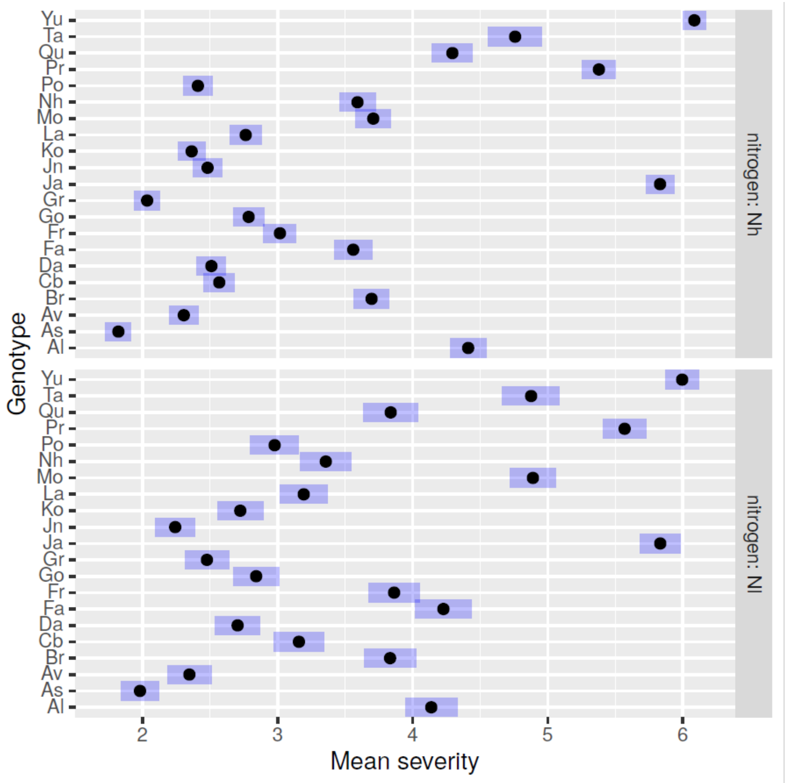

**S-3. Summary figures for each of the three datasets and pairwise comparisons of least square means across Severity classes and across Genotypes**

**Fig. S3.1** Graphical examination of the fraction (F/S) of fruiting bodies pixels in states F (fruiting bodies) and S (stem) on 3029 oilseed rape stems after the processing and post-processing of RGB digital images in the Data1416 dataset. a) Histogram of class frequency depending on fruiting bodies fraction ; b-d) , Boxplots showing how the fraction F/S changed b) between 2 Years; c) between 7 classes of phoma stem canker Severity before incubation; d) between 15 Genotypes. Phoma stem canker severity prior incubation classes S1 to S6 correspond to increasing proportion of cankered cross section at crown level. Year, disease Severity and Genotype all have a significant effect.

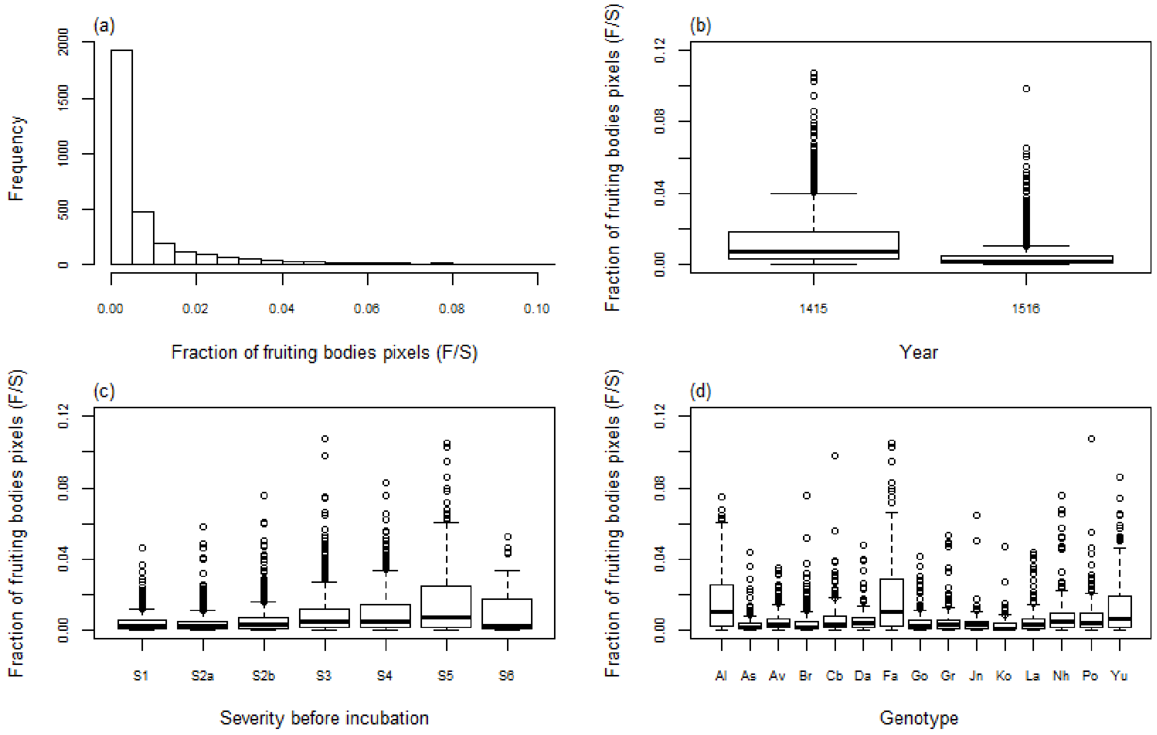

**Table S3.1** The pairwise comparisons of least-square means pointed out a significant difference between the Severity classes in the Data1416 dataset. Results are averaged over the levels of severity for each of the 2 Years and the 2 Nitrogen fertilization levels; Confidence level used: 0.95; P value adjustment: Tukey method for comparing a family of 7 estimates; Significance level used: alpha = 0.05.

| Year | Nitrogen | Severity | Prob. | SE | df | asymp.LCL | asymp.UCL | .group |
|---|---|---|---|---|---|---|---|---|
| 1415 | Nh | S1 | 0,007 | 0,000 | Inf | 0,007 | 0,007 | 1 |
| 1415 | Nh | S2a | 0,008 | 0,000 | Inf | 0,008 | 0,008 | 2 |
| 1415 | Nh | S2b | 0,012 | 0,000 | Inf | 0,012 | 0,012 | 3 |
| 1415 | Nh | S4 | 0,018 | 0,000 | Inf | 0,018 | 0,018 | 4 |
| 1415 | Nh | S3 | 0,020 | 0,000 | Inf | 0,020 | 0,020 | 5 |
| 1415 | Nh | S6 | 0,022 | 0,000 | Inf | 0,022 | 0,022 | 6 |
| 1415 | Nh | S5 | 0,029 | 0,000 | Inf | 0,029 | 0,029 | 7 |
| 1415 | Nl | S1 | 0,005 | 0,000 | Inf | 0,005 | 0,005 | 1 |
| 1415 | Nl | S2a | 0,006 | 0,000 | Inf | 0,006 | 0,006 | 2 |
| 1415 | Nl | S2b | 0,008 | 0,000 | Inf | 0,008 | 0,008 | 3 |
| 1415 | Nl | S3 | 0,015 | 0,000 | Inf | 0,015 | 0,015 | 4 |
| 1415 | Nl | S6 | 0,018 | 0,000 | Inf | 0,018 | 0,018 | 5 |
| 1415 | Nl | S5 | 0,019 | 0,000 | Inf | 0,019 | 0,019 | 6 |
| 1415 | Nl | S4 | 0,019 | 0,000 | Inf | 0,019 | 0,019 | 6 |
| 1516 | Nh | S1 | 0,003 | 0,000 | Inf | 0,003 | 0,003 | 1 |
| 1516 | Nh | S2a | 0,003 | 0,000 | Inf | 0,003 | 0,003 | 1 |
| 1516 | Nh | S2b | 0,005 | 0,000 | Inf | 0,005 | 0,005 | 2 |
| 1516 | Nh | S3 | 0,005 | 0,000 | Inf | 0,005 | 0,005 | 3 |
| 1516 | Nh | S4 | 0,005 | 0,000 | Inf | 0,005 | 0,006 | 4 |
| 1516 | Nh | S6 | 0,006 | 0,000 | Inf | 0,006 | 0,006 | 4 |
| 1516 | Nh | S5 | 0,009 | 0,000 | Inf | 0,009 | 0,009 | 5 |
| 1516 | Nl | S6 | 0,003 | 0,000 | Inf | 0,003 | 0,003 | 1 |
| 1516 | Nl | S2a | 0,003 | 0,000 | Inf | 0,003 | 0,003 | 2 |
| 1516 | Nl | S2b | 0,003 | 0,000 | Inf | 0,003 | 0,003 | 3 |
| 1516 | Nl | S1 | 0,003 | 0,000 | Inf | 0,003 | 0,003 | 4 |
| 1516 | Nl | S3 | 0,004 | 0,000 | Inf | 0,004 | 0,004 | 5 |
| 1516 | Nl | S5 | 0,005 | 0,000 | Inf | 0,005 | 0,005 | 6 |
| 1516 | Nl | S4 | 0,006 | 0,000 | Inf | 0,006 | 0,006 | 7 |

**Figure S3.2** Predicted probability of fruiting bodies pixels by Severity class for each of the 2 Years and the 2 Nitrogen fertilization levels in the Data1416 dataset.

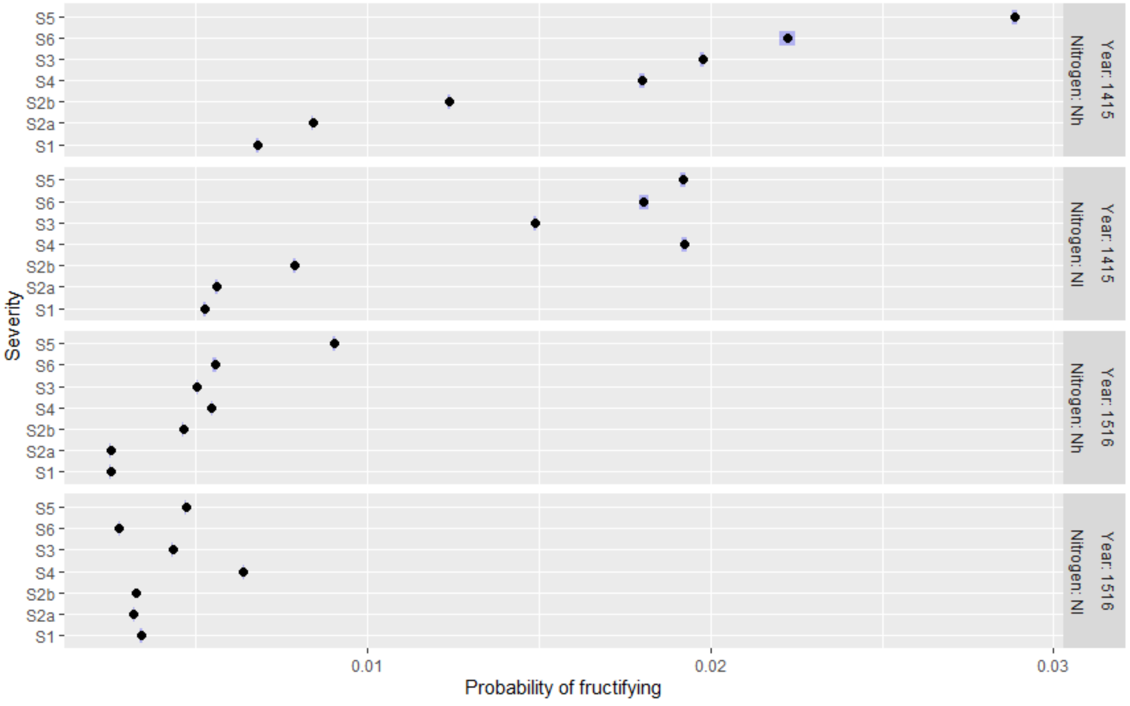

**Table S3.2** The pairwise comparisons of least-square means pointed out a significant difference between the genotypes in the Data1416 dataset. Results are averaged over the levels of Genotypes for each of the 2 Years and the 2 Nitrogen fertilization levels; Confidence level used: 0.95; P value adjustment: Tukey method for comparing a family of 15 estimates; Significance level used: alpha = 0.05.

| Year | Nitrogen | Genotype | prob | SE | df | asymp.LCL | asymp.UCL | group |
|---|---|---|---|---|---|---|---|---|
| 1415 | Nh | Ko | 0,008 | 0,000 | Inf | 0,008 | 0,008 | 1 |
| 1415 | Nh | Go | 0,008 | 0,000 | Inf | 0,008 | 0,009 | 1 |
| 1415 | Nh | As | 0,010 | 0,000 | Inf | 0,010 | 0,010 | 2 |
| 1415 | Nh | Cb | 0,012 | 0,000 | Inf | 0,012 | 0,012 | 3 |
| 1415 | Nh | Jn | 0,012 | 0,000 | Inf | 0,012 | 0,012 | 3 |
| 1415 | Nh | Av | 0,014 | 0,000 | Inf | 0,014 | 0,014 | 4 |
| 1415 | Nh | Da | 0,014 | 0,000 | Inf | 0,014 | 0,014 | 5 |
| 1415 | Nh | Nh | 0,015 | 0,000 | Inf | 0,015 | 0,015 | 6 |
| 1415 | Nh | La | 0,015 | 0,000 | Inf | 0,015 | 0,015 | 6 |
| 1415 | Nh | Br | 0,018 | 0,000 | Inf | 0,018 | 0,018 | 7 |
| 1415 | Nh | Yu | 0,019 | 0,000 | Inf | 0,018 | 0,019 | 8 |
| 1415 | Nh | Gr | 0,019 | 0,000 | Inf | 0,019 | 0,019 | 9 |
| 1415 | Nh | Po | 0,024 | 0,000 | Inf | 0,023 | 0,024 | 0 |
| 1415 | Nh | Al | 0,027 | 0,000 | Inf | 0,027 | 0,027 | A |
| 1415 | Nh | Fa | 0,034 | 0,000 | Inf | 0,034 | 0,034 | B |
| 1415 | Nl | Ko | 0,007 | 0,000 | Inf | 0,007 | 0,007 | 1 |
| 1415 | Nl | As | 0,008 | 0,000 | Inf | 0,008 | 0,008 | 2 |
| 1415 | Nl | Br | 0,009 | 0,000 | Inf | 0,009 | 0,009 | 3 |
| 1415 | Nl | Jn | 0,011 | 0,000 | Inf | 0,011 | 0,011 | 4 |
| 1415 | Nl | Gr | 0,011 | 0,000 | Inf | 0,011 | 0,011 | 5 |
| 1415 | Nl | Go | 0,012 | 0,000 | Inf | 0,011 | 0,012 | 6 |
| 1415 | Nl | Da | 0,013 | 0,000 | Inf | 0,013 | 0,013 | 7 |
| 1415 | Nl | La | 0,013 | 0,000 | Inf | 0,013 | 0,013 | 7 |
| 1415 | Nl | Nh | 0,013 | 0,000 | Inf | 0,013 | 0,013 | 8 |
| 1415 | Nl | Yu | 0,014 | 0,000 | Inf | 0,014 | 0,014 | 9 |
| 1415 | Nl | Av | 0,014 | 0,000 | Inf | 0,014 | 0,014 | 0 |
| 1415 | Nl | Po | 0,015 | 0,000 | Inf | 0,015 | 0,015 | A |
| 1415 | Nl | Cb | 0,017 | 0,000 | Inf | 0,017 | 0,017 | B |
| 1415 | Nl | Al | 0,017 | 0,000 | Inf | 0,017 | 0,017 | B |
| 1415 | Nl | Fa | 0,018 | 0,000 | Inf | 0,018 | 0,018 | C |
| 1516 | Nh | Br | 0,002 | 0,000 | Inf | 0,002 | 0,002 | 1 |
| 1516 | Nh | La | 0,003 | 0,000 | Inf | 0,003 | 0,003 | 2 |
| 1516 | Nh | Ko | 0,004 | 0,000 | Inf | 0,004 | 0,004 | 3 |
| 1516 | Nh | Gr | 0,004 | 0,000 | Inf | 0,004 | 0,004 | 4 |
| 1516 | Nh | Nh | 0,004 | 0,000 | Inf | 0,004 | 0,004 | 5 |
| 1516 | Nh | Po | 0,004 | 0,000 | Inf | 0,004 | 0,004 | 6 |
| 1516 | Nh | Jn | 0,004 | 0,000 | Inf | 0,004 | 0,004 | 6 |
| 1516 | Nh | Yu | 0,004 | 0,000 | Inf | 0,004 | 0,004 | 6 |
| 1516 | Nh | As | 0,004 | 0,000 | Inf | 0,004 | 0,004 | 7 |
| 1516 | Nh | Av | 0,006 | 0,000 | Inf | 0,006 | 0,006 | 8 |
| 1516 | Nh | Go | 0,006 | 0,000 | Inf | 0,006 | 0,006 | 9 |
| 1516 | Nh | Da | 0,007 | 0,000 | Inf | 0,007 | 0,007 | 0 |
| 1516 | Nh | Al | 0,007 | 0,000 | Inf | 0,007 | 0,007 | A |
| 1516 | Nh | Cb | 0,007 | 0,000 | Inf | 0,007 | 0,007 | B |
| 1516 | Nh | Fa | 0,008 | 0,000 | Inf | 0,008 | 0,008 | C |
| 1516 | Nl | Ko | 0,001 | 0,000 | Inf | 0,001 | 0,001 | 1 |
| 1516 | Nl | Br | 0,002 | 0,000 | Inf | 0,002 | 0,002 | 2 |
| 1516 | Nl | Jn | 0,002 | 0,000 | Inf | 0,002 | 0,002 | 3 |
| 1516 | Nl | Yu | 0,002 | 0,000 | Inf | 0,002 | 0,002 | 4 |
| 1516 | Nl | As | 0,002 | 0,000 | Inf | 0,002 | 0,002 | 45 |
| 1516 | Nl | La | 0,002 | 0,000 | Inf | 0,002 | 0,002 | 5 |
| 1516 | Nl | Nh | 0,003 | 0,000 | Inf | 0,003 | 0,003 | 6 |
| 1516 | Nl | Go | 0,003 | 0,000 | Inf | 0,003 | 0,003 | 7 |
| 1516 | Nl | Da | 0,003 | 0,000 | Inf | 0,003 | 0,003 | 8 |
| 1516 | Nl | Gr | 0,003 | 0,000 | Inf | 0,003 | 0,003 | 9 |
| 1516 | Nl | Cb | 0,004 | 0,000 | Inf | 0,004 | 0,004 | 0 |
| 1516 | Nl | Av | 0,004 | 0,000 | Inf | 0,004 | 0,004 | A |
| 1516 | Nl | Po | 0,005 | 0,000 | Inf | 0,005 | 0,005 | B |
| 1516 | Nl | Al | 0,006 | 0,000 | Inf | 0,006 | 0,006 | C |
| 1516 | Nl | Fa | 0,017 | 0,000 | Inf | 0,017 | 0,018 | D |

**Fig. S3.3** Predicted probability of fruiting bodies pixels by Genotype for each of the 2 Years and the 2 Nitrogen fertilization levels in the Data1416 dataset.

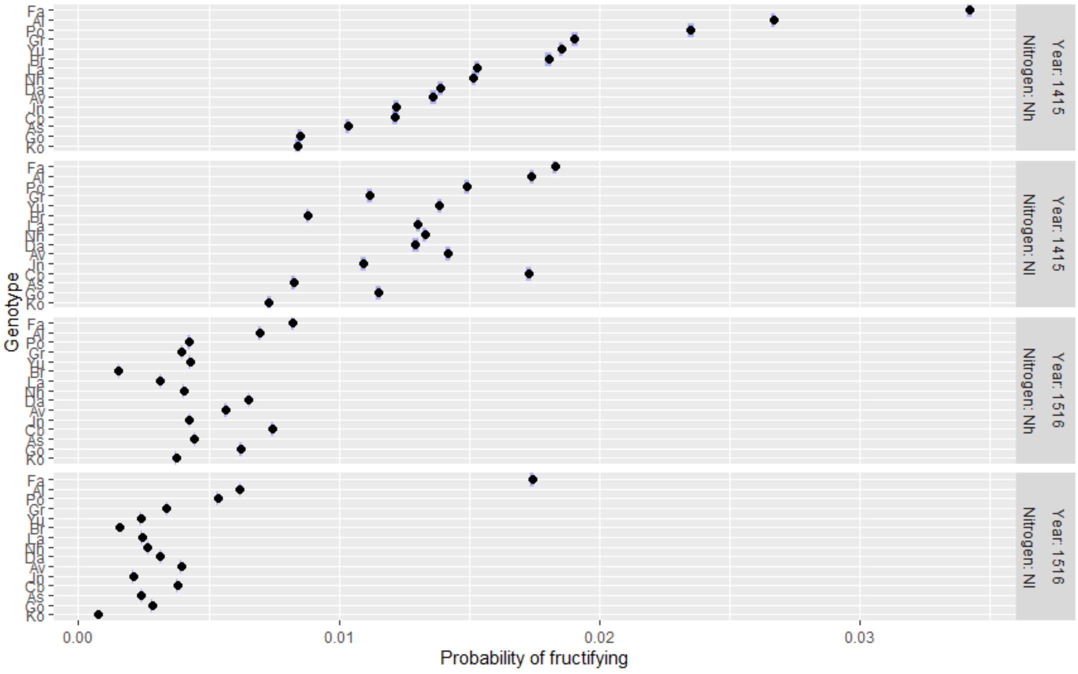

**Fig. S3.4** Graphical examination of the fraction (F/S) of fruiting bodies pixels in states F (fruiting bodies) and S (stem) on 3379 oilseed rape stems after the processing and post-processing of RGB digital images in the Data1216 dataset. a) Histogram of class frequency depending on fruiting bodies fraction ; b-d) , Boxplots showing how the fraction F/S changed b) between 4 Years; c) between 7 classes of phoma stem canker Severity before incubation; d) between 15 Genotypes. Phoma stem canker severity prior incubation classes S1 to S6 correspond to increasing proportion of cankered cross section at crown level. Year, disease Severity and Genotype all have a significant effect.

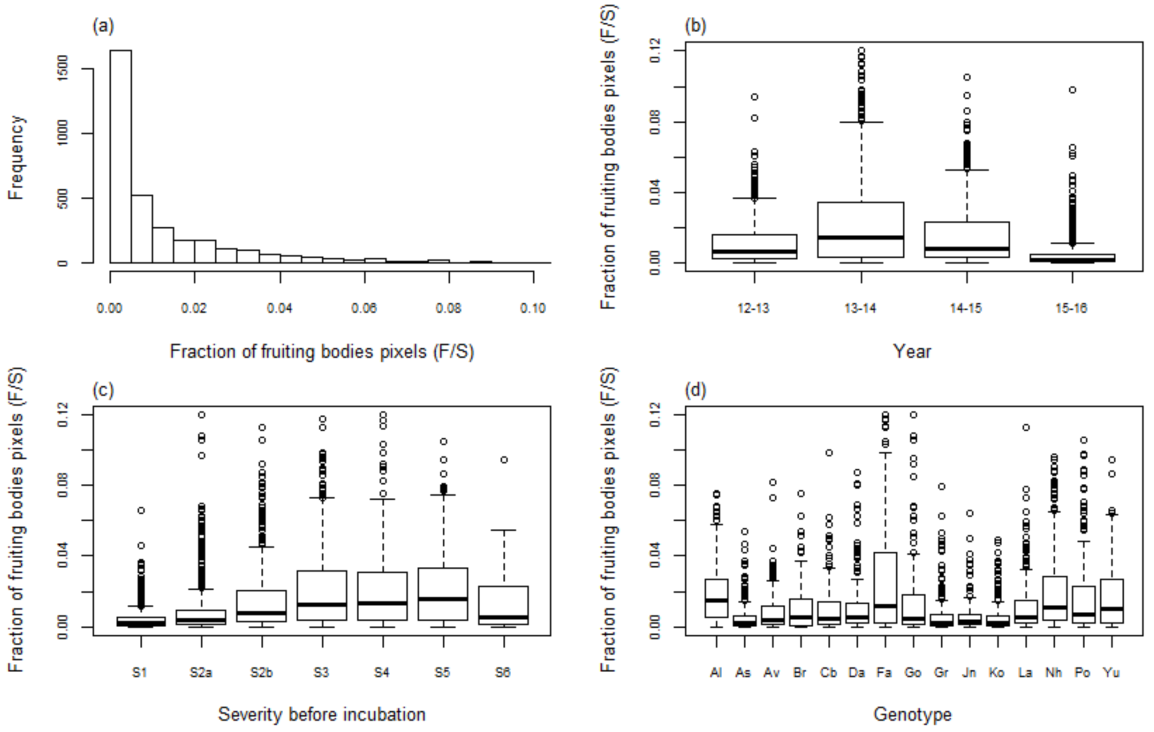

**Table S3.3** The pairwise comparisons of least-square means pointed out a significant difference between the Severity classes in the Data1216 dataset. Results are averaged over the levels of severity for each of the 4 Years and the Nh Nitrogen fertilization level; Confidence level used: 0.95; P value adjustment: Tukey method for comparing a family of 7 estimates; Significance level used: alpha = 0.05.

| Year | Severity | Prob. | SE | df | asymp.LCL | asymp.UCL | Group |
|---|---|---|---|---|---|---|---|
| 1213 | S1 | 0,003 | 0,000 | Inf | 0,003 | 0,003 | 1 |
| 1213 | S2a | 0,010 | 0,000 | Inf | 0,010 | 0,010 | 2 |
| 1213 | S2b | 0,015 | 0,000 | Inf | 0,015 | 0,015 | 3 |
| 1213 | S3 | 0,016 | 0,000 | Inf | 0,016 | 0,016 | 4 |
| 1213 | S5 | 0,020 | 0,000 | Inf | 0,020 | 0,020 | 5 |
| 1213 | S4 | 0,021 | 0,000 | Inf | 0,020 | 0,021 | 6 |
| 1213 | S6 | 0,024 | 0,000 | Inf | 0,024 | 0,024 | 7 |
| 1314 | S1 | 0,008 | 0,000 | Inf | 0,008 | 0,008 | 1 |
| 1314 | S6 | 0,012 | 0,000 | Inf | 0,012 | 0,012 | 2 |
| 1314 | S2a | 0,013 | 0,000 | Inf | 0,013 | 0,013 | 3 |
| 1314 | S2b | 0,024 | 0,000 | Inf | 0,024 | 0,024 | 4 |
| 1314 | S4 | 0,028 | 0,000 | Inf | 0,028 | 0,028 | 5 |
| 1314 | S5 | 0,029 | 0,000 | Inf | 0,029 | 0,029 | 6 |
| 1314 | S3 | 0,032 | 0,000 | Inf | 0,032 | 0,032 | 7 |
| 1415 | S1 | 0,007 | 0,000 | Inf | 0,007 | 0,007 | 1 |
| 1415 | S2a | 0,008 | 0,000 | Inf | 0,008 | 0,008 | 2 |
| 1415 | S2b | 0,012 | 0,000 | Inf | 0,012 | 0,012 | 3 |
| 1415 | S4 | 0,018 | 0,000 | Inf | 0,018 | 0,018 | 4 |
| 1415 | S3 | 0,020 | 0,000 | Inf | 0,020 | 0,020 | 5 |
| 1415 | S6 | 0,022 | 0,000 | Inf | 0,022 | 0,022 | 6 |
| 1415 | S5 | 0,029 | 0,000 | Inf | 0,029 | 0,029 | 7 |
| 1516 | S1 | 0,003 | 0,000 | Inf | 0,003 | 0,003 | 1 |
| 1516 | S2a | 0,003 | 0,000 | Inf | 0,003 | 0,003 | 1 |
| 1516 | S2b | 0,005 | 0,000 | Inf | 0,005 | 0,005 | 2 |
| 1516 | S3 | 0,005 | 0,000 | Inf | 0,005 | 0,005 | 3 |
| 1516 | S4 | 0,005 | 0,000 | Inf | 0,005 | 0,006 | 4 |
| 1516 | S6 | 0,006 | 0,000 | Inf | 0,006 | 0,006 | 4 |
| 1516 | S5 | 0,009 | 0,000 | Inf | 0,009 | 0,009 | 5 |

**Fig. S3.5** Predicted probability of fruiting bodies pixels by Severity class for each of the 4 Years and the Nh Nitrogen fertilization level in the Data1216 dataset.

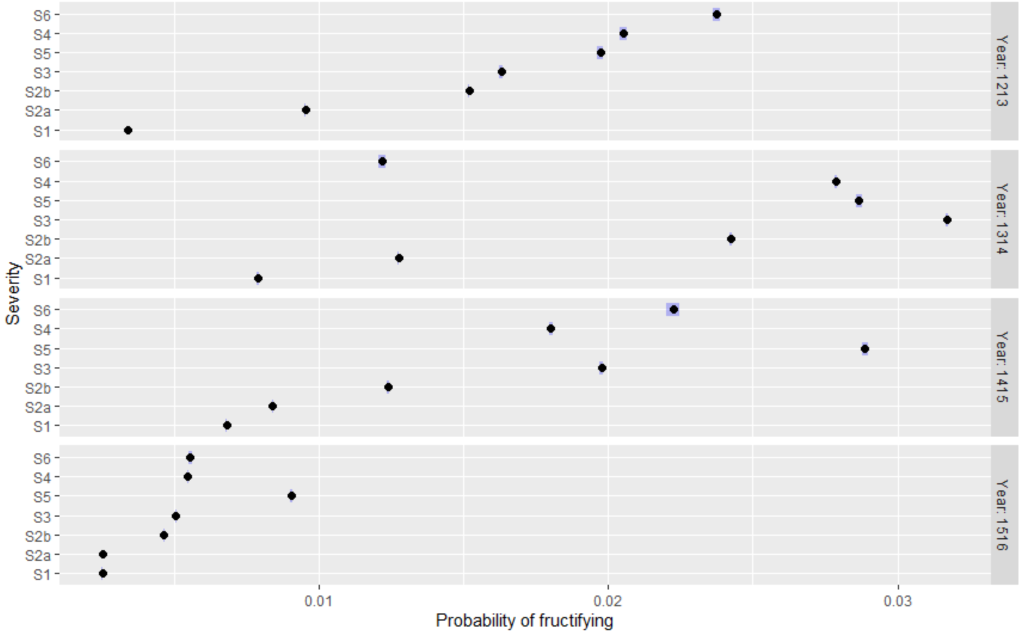

**Table S3.4** The pairwise comparisons of least-square means pointed out a significant difference between the genotypes in the Data1216 dataset. Results are averaged over the levels of Genotypes for each of the 4 Years and the Nh Nitrogen fertilization level; Confidence level used: 0.95; P value adjustment: Tukey method for comparing a family of 15 estimates; Significance level used: alpha = 0.05.

| Year | Genotype | prob | SE | df | asymp.LCL | asymp.UCL | group |
|---|---|---|---|---|---|---|---|
| 1213 | As | 0,009 | 0,000 | Inf | 0,009 | 0,009 | 1 |
| 1213 | La | 0,009 | 0,000 | Inf | 0,009 | 0,010 | 2 |
| 1213 | Cb | 0,011 | 0,000 | Inf | 0,011 | 0,011 | 3 |
| 1213 | Fa | 0,011 | 0,000 | Inf | 0,011 | 0,012 | 4 |
| 1213 | Ko | 0,012 | 0,000 | Inf | 0,012 | 0,013 | 5 |
| 1213 | Al | 0,013 | 0,000 | Inf | 0,013 | 0,013 | 6 |
| 1213 | Jn | 0,013 | 0,000 | Inf | 0,013 | 0,013 | 6 |
| 1213 | Yu | 0,016 | 0,000 | Inf | 0,016 | 0,016 | 7 |
| 1213 | Da | 0,017 | 0,000 | Inf | 0,016 | 0,017 | 8 |
| 1213 | Gr | 0,018 | 0,000 | Inf | 0,017 | 0,018 | 9 |
| 1213 | Br | 0,018 | 0,000 | Inf | 0,018 | 0,018 | 0 |
| 1213 | Nh | 0,020 | 0,000 | Inf | 0,020 | 0,020 | A |
| 1213 | Go | 0,021 | 0,000 | Inf | 0,021 | 0,021 | B |
| 1213 | Po | 0,023 | 0,000 | Inf | 0,023 | 0,023 | C |
| 1213 | Av | 0,023 | 0,000 | Inf | 0,023 | 0,023 | D |
| 1314 | Jn | 0,008 | 0,000 | Inf | 0,007 | 0,008 | 1 |
| 1314 | Av | 0,009 | 0,000 | Inf | 0,009 | 0,009 | 2 |
| 1314 | Gr | 0,012 | 0,000 | Inf | 0,012 | 0,012 | 3 |
| 1314 | Ko | 0,012 | 0,000 | Inf | 0,012 | 0,013 | 4 |
| 1314 | Br | 0,013 | 0,000 | Inf | 0,013 | 0,013 | 5 |
| 1314 | As | 0,014 | 0,000 | Inf | 0,014 | 0,014 | 6 |
| 1314 | Yu | 0,015 | 0,000 | Inf | 0,015 | 0,015 | 7 |
| 1314 | Cb | 0,018 | 0,000 | Inf | 0,018 | 0,018 | 8 |
| 1314 | Al | 0,019 | 0,000 | Inf | 0,019 | 0,019 | 9 |
| 1314 | La | 0,023 | 0,000 | Inf | 0,023 | 0,023 | 0 |
| 1314 | Da | 0,026 | 0,000 | Inf | 0,026 | 0,026 | A |
| 1314 | Po | 0,030 | 0,000 | Inf | 0,030 | 0,030 | B |
| 1314 | Nh | 0,033 | 0,000 | Inf | 0,033 | 0,033 | C |
| 1314 | Go | 0,038 | 0,000 | Inf | 0,038 | 0,038 | D |
| 1314 | Fa | 0,042 | 0,000 | Inf | 0,042 | 0,042 | E |
| 1415 | Ko | 0,008 | 0,000 | Inf | 0,008 | 0,008 | 1 |
| 1415 | Go | 0,008 | 0,000 | Inf | 0,008 | 0,009 | 1 |
| 1415 | As | 0,010 | 0,000 | Inf | 0,010 | 0,010 | 2 |
| 1415 | Cb | 0,012 | 0,000 | Inf | 0,012 | 0,012 | 3 |
| 1415 | Jn | 0,012 | 0,000 | Inf | 0,012 | 0,012 | 3 |
| 1415 | Av | 0,014 | 0,000 | Inf | 0,014 | 0,014 | 4 |
| 1415 | Da | 0,014 | 0,000 | Inf | 0,014 | 0,014 | 5 |
| 1415 | Nh | 0,015 | 0,000 | Inf | 0,015 | 0,015 | 6 |
| 1415 | La | 0,015 | 0,000 | Inf | 0,015 | 0,015 | 6 |
| 1415 | Br | 0,018 | 0,000 | Inf | 0,018 | 0,018 | 7 |
| 1415 | Yu | 0,019 | 0,000 | Inf | 0,018 | 0,019 | 8 |
| 1415 | Gr | 0,019 | 0,000 | Inf | 0,019 | 0,019 | 9 |
| 1415 | Po | 0,024 | 0,000 | Inf | 0,023 | 0,024 | 0 |
| 1415 | Al | 0,027 | 0,000 | Inf | 0,027 | 0,027 | A |
| 1415 | Fa | 0,034 | 0,000 | Inf | 0,034 | 0,034 | B |
| 1516 | Br | 0,002 | 0,000 | Inf | 0,002 | 0,002 | 1 |
| 1516 | La | 0,003 | 0,000 | Inf | 0,003 | 0,003 | 2 |
| 1516 | Ko | 0,004 | 0,000 | Inf | 0,004 | 0,004 | 3 |
| 1516 | Gr | 0,004 | 0,000 | Inf | 0,004 | 0,004 | 4 |
| 1516 | Nh | 0,004 | 0,000 | Inf | 0,004 | 0,004 | 5 |
| 1516 | Po | 0,004 | 0,000 | Inf | 0,004 | 0,004 | 6 |
| 1516 | Jn | 0,004 | 0,000 | Inf | 0,004 | 0,004 | 6 |
| 1516 | Yu | 0,004 | 0,000 | Inf | 0,004 | 0,004 | 6 |
| 1516 | As | 0,004 | 0,000 | Inf | 0,004 | 0,004 | 7 |
| 1516 | Av | 0,006 | 0,000 | Inf | 0,006 | 0,006 | 8 |
| 1516 | Go | 0,006 | 0,000 | Inf | 0,006 | 0,006 | 9 |
| 1516 | Da | 0,007 | 0,000 | Inf | 0,007 | 0,007 | 0 |
| 1516 | Al | 0,007 | 0,000 | Inf | 0,007 | 0,007 | A |
| 1516 | Cb | 0,007 | 0,000 | Inf | 0,007 | 0,007 | B |
| 1516 | Fa | 0,008 | 0,000 | Inf | 0,008 | 0,008 | C |

**Fig. S3.6** Predicted probability of fruiting bodies pixels by Genotype for each of the 4 Years and the Nh Nitrogen fertilization level in the Data1416 dataset.

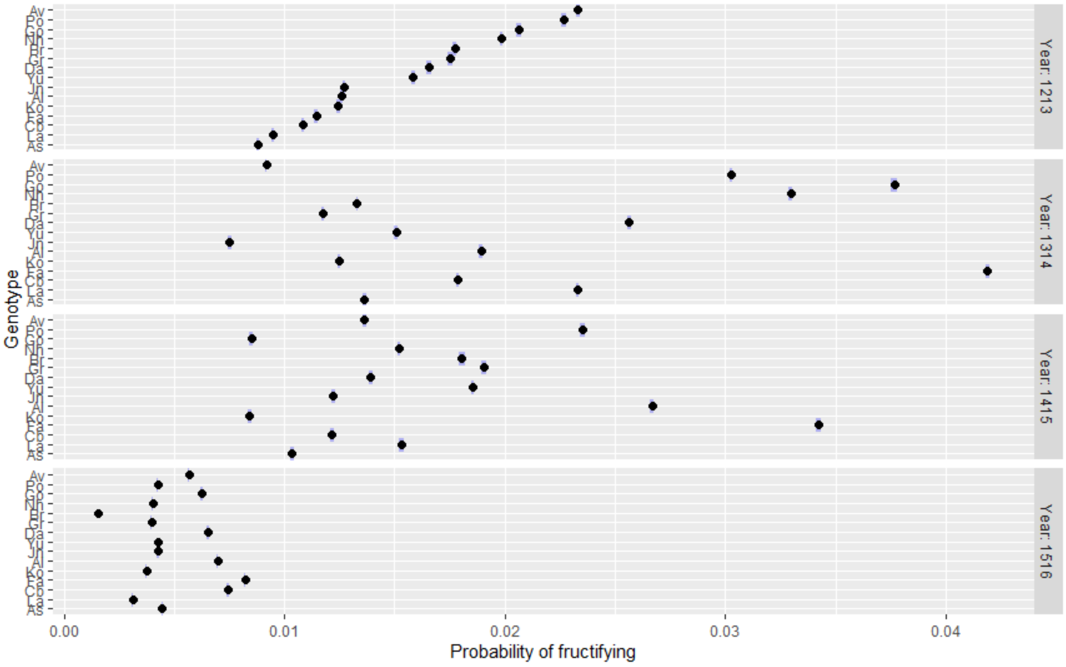

**Fig. S3.7** Graphical examination of the fraction (F/S) of fruiting bodies pixels in states F (fruiting bodies) and S (stem) on 2677 oilseed rape stems after the processing and post-processing of RGB digital images in the Data1516 dataset. a) Histogram of class frequency depending on fruiting bodies fraction ; b-d) , Boxplots showing how the fraction F/S changed b) between 4 Years; c) between 7 classes of phoma stem canker Severity before incubation; d) between 21 Genotypes. Phoma stem canker severity prior incubation classes S1 to S6 correspond to increasing proportion of cankered cross section at crown level. Disease Severity, and Genotype all have a significant effect.

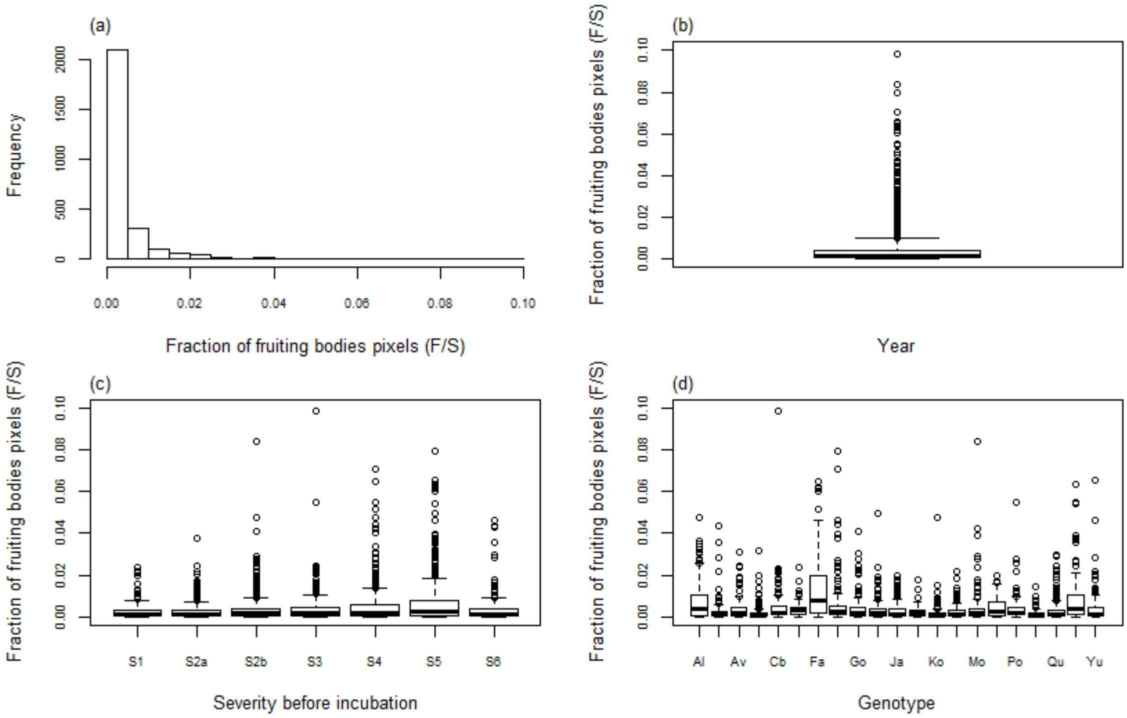

**Table S3.5** The pairwise comparisons of least-square means pointed out a significant difference between the Severity classes in the Data1516 dataset. Results are averaged over the levels of severity for the 1516 Year and the 2 Nitrogen fertilization level; Confidence level used: 0.95; P value adjustment: Tukey method for comparing a family of 7 estimates; Significance level used: alpha = 0.05.

| Severity | Nitrogen | prob | SE | df | asymp.LCL | asymp.UCL | group |
|---|---|---|---|---|---|---|---|
| S1 | Nh | 0,002 | 0,000 | Inf | 0,002 | 0,002 | 1 |
| S2a | Nh | 0,002 | 0,000 | Inf | 0,002 | 0,002 | 1 |
| S2b | Nh | 0,004 | 0,000 | Inf | 0,004 | 0,004 | 2 |
| S3 | Nh | 0,004 | 0,000 | Inf | 0,004 | 0,005 | 3 |
| S4 | Nh | 0,005 | 0,000 | Inf | 0,005 | 0,005 | 4 |
| S6 | Nh | 0,006 | 0,000 | Inf | 0,006 | 0,006 | 5 |
| S5 | Nh | 0,009 | 0,000 | Inf | 0,009 | 0,009 | 6 |
| S6 | Nl | 0,003 | 0,000 | Inf | 0,003 | 0,003 | 1 |
| S1 | Nl | 0,003 | 0,000 | Inf | 0,003 | 0,003 | 2 |
| S2a | Nl | 0,003 | 0,000 | Inf | 0,003 | 0,003 | 3 |
| S2b | Nl | 0,004 | 0,000 | Inf | 0,004 | 0,004 | 4 |
| S3 | Nl | 0,004 | 0,000 | Inf | 0,004 | 0,004 | 5 |
| S4 | Nl | 0,006 | 0,000 | Inf | 0,006 | 0,006 | 6 |
| S5 | Nl | 0,006 | 0,000 | Inf | 0,006 | 0,006 | 7 |

**Fig. S3.8** Predicted probability of fruiting bodies pixels by Severity class for the 1516 Year and the 2 Nitrogen fertilization level in the Data1516 dataset.

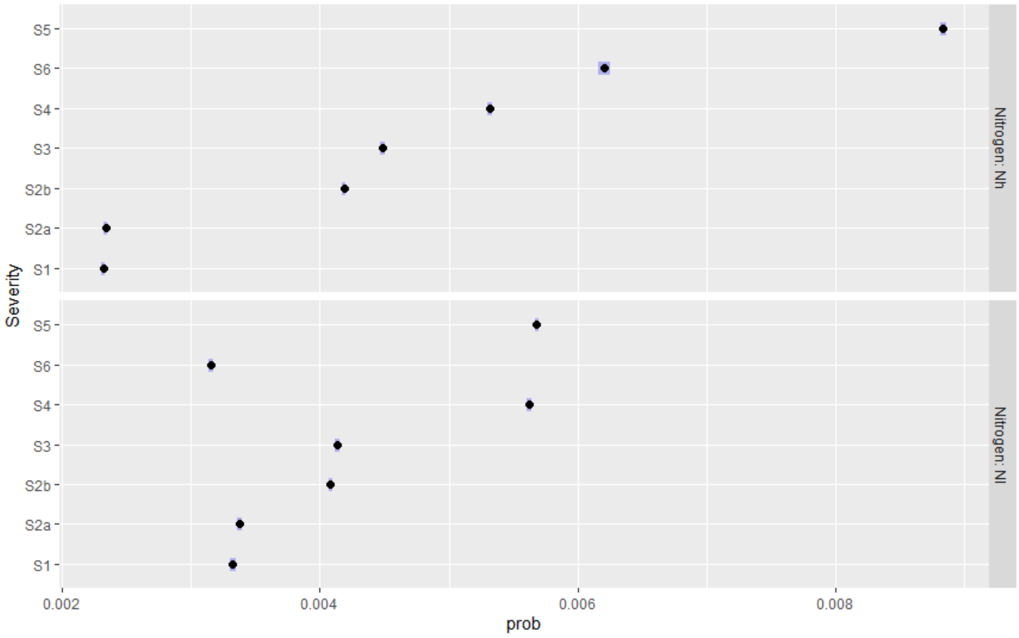

**Table S3.6** The pairwise comparisons of least-square means pointed out a significant difference between the genotypes in the Data1516 dataset. Results are averaged over the levels of Genotypes for the 1516 Year and the 2 Nitrogen fertilization level; Confidence level used: 0.95; P value adjustment: Tukey method for comparing a family of 21 estimates; Significance level used alpha = 0.05.

| Genotype | Nitrogen | prob | SE | df | asymp.LCL | asymp.UCL | group |
|---|---|---|---|---|---|---|---|
| Pr | Nh | 0,001 | 0,000 | Inf | 0,001 | 0,001 | 1 |
| Br | Nh | 0,002 | 0,000 | Inf | 0,002 | 0,002 | 2 |
| Mo | Nh | 0,002 | 0,000 | Inf | 0,002 | 0,002 | 3 |
| Ja | Nh | 0,002 | 0,000 | Inf | 0,002 | 0,002 | 4 |
| Qu | Nh | 0,003 | 0,000 | Inf | 0,003 | 0,003 | 5 |
| La | Nh | 0,003 | 0,000 | Inf | 0,003 | 0,003 | 6 |
| Ko | Nh | 0,004 | 0,000 | Inf | 0,004 | 0,004 | 7 |
| Yu | Nh | 0,004 | 0,000 | Inf | 0,004 | 0,004 | 7 |
| Gr | Nh | 0,004 | 0,000 | Inf | 0,004 | 0,004 | 8 |
| Nh | Nh | 0,004 | 0,000 | Inf | 0,004 | 0,004 | 8 |
| Jn | Nh | 0,004 | 0,000 | Inf | 0,004 | 0,005 | 9 |
| Po | Nh | 0,005 | 0,000 | Inf | 0,004 | 0,005 | 9 |
| As | Nh | 0,005 | 0,000 | Inf | 0,005 | 0,005 | 9 |
| Ta | Nh | 0,005 | 0,000 | Inf | 0,005 | 0,006 | 0 |
| Av | Nh | 0,006 | 0,000 | Inf | 0,006 | 0,006 | A |
| Go | Nh | 0,007 | 0,000 | Inf | 0,007 | 0,007 | B |
| Da | Nh | 0,007 | 0,000 | Inf | 0,007 | 0,007 | C |
| Al | Nh | 0,007 | 0,000 | Inf | 0,007 | 0,007 | D |
| Cb | Nh | 0,008 | 0,000 | Inf | 0,008 | 0,008 | E |
| Fa | Nh | 0,008 | 0,000 | Inf | 0,008 | 0,008 | F |
| Fr | Nh | 0,010 | 0,000 | Inf | 0,010 | 0,010 | G |
| Ko | Nl | 0,001 | 0,000 | Inf | 0,001 | 0,001 | 1 |
| Pr | Nl | 0,001 | 0,000 | Inf | 0,001 | 0,001 | 2 |
| Br | Nl | 0,002 | 0,000 | Inf | 0,002 | 0,002 | 3 |
| Jn | Nl | 0,002 | 0,000 | Inf | 0,002 | 0,002 | 4 |
| Ja | Nl | 0,002 | 0,000 | Inf | 0,002 | 0,002 | 5 |
| Yu | Nl | 0,002 | 0,000 | Inf | 0,002 | 0,002 | 5 |
| As | Nl | 0,002 | 0,000 | Inf | 0,002 | 0,002 | 6 |
| La | Nl | 0,002 | 0,000 | Inf | 0,002 | 0,003 | 7 |
| Nh | Nl | 0,003 | 0,000 | Inf | 0,003 | 0,003 | 8 |
| Go | Nl | 0,003 | 0,000 | Inf | 0,003 | 0,003 | 9 |
| Da | Nl | 0,003 | 0,000 | Inf | 0,003 | 0,003 | 0 |
| Gr | Nl | 0,003 | 0,000 | Inf | 0,003 | 0,003 | A |
| Fr | Nl | 0,003 | 0,000 | Inf | 0,003 | 0,003 | B |
| Qu | Nl | 0,003 | 0,000 | Inf | 0,003 | 0,003 | B |
| Cb | Nl | 0,004 | 0,000 | Inf | 0,004 | 0,004 | C |
| Av | Nl | 0,004 | 0,000 | Inf | 0,004 | 0,004 | D |
| Po | Nl | 0,005 | 0,000 | Inf | 0,005 | 0,005 | E |
| Al | Nl | 0,006 | 0,000 | Inf | 0,006 | 0,006 | F |
| Mo | Nl | 0,007 | 0,000 | Inf | 0,007 | 0,007 | G |
| Ta | Nl | 0,011 | 0,000 | Inf | 0,011 | 0,011 | H |
| Fa | Nl | 0,016 | 0,000 | Inf | 0,016 | 0,017 | I |

**Fig. S3.9** Predicted probability of fruiting bodies pixels by Genotype for the 1516 Year and the 2 Nitrogen fertilization level in the Data1516 dataset.

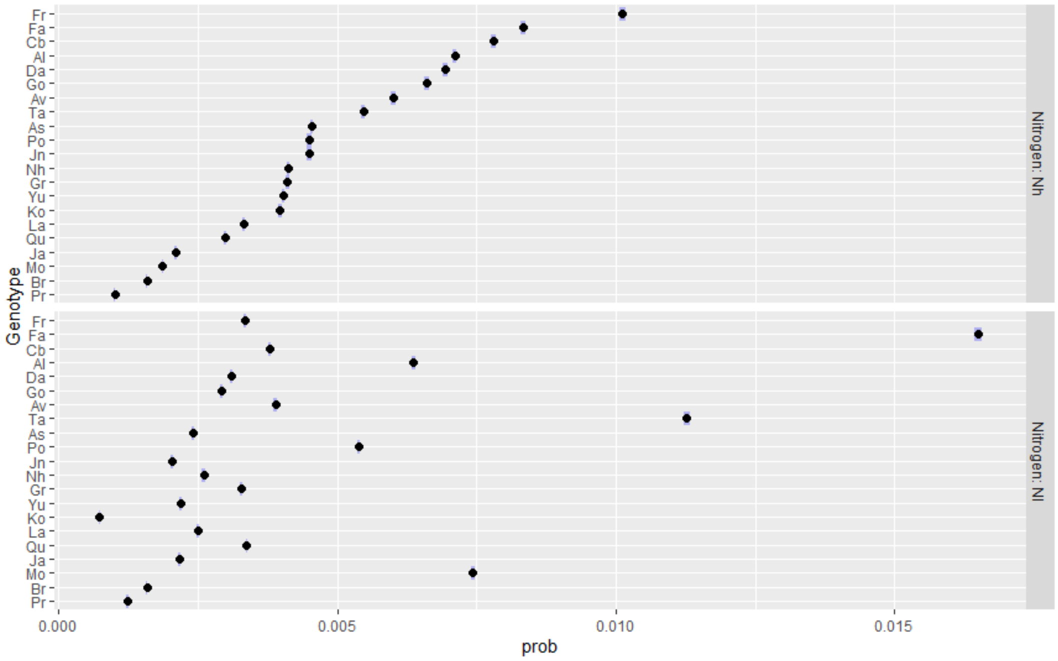

**S-4. Fraction of fruiting bodies pixels depending on the severity before incubation for each of the genotypes in each of the datasets.**

**Fig. S4.1.** Fraction of fruiting bodies pixels depending on the severity before incubation for the 15 genotypes in the Data1216 dataset.

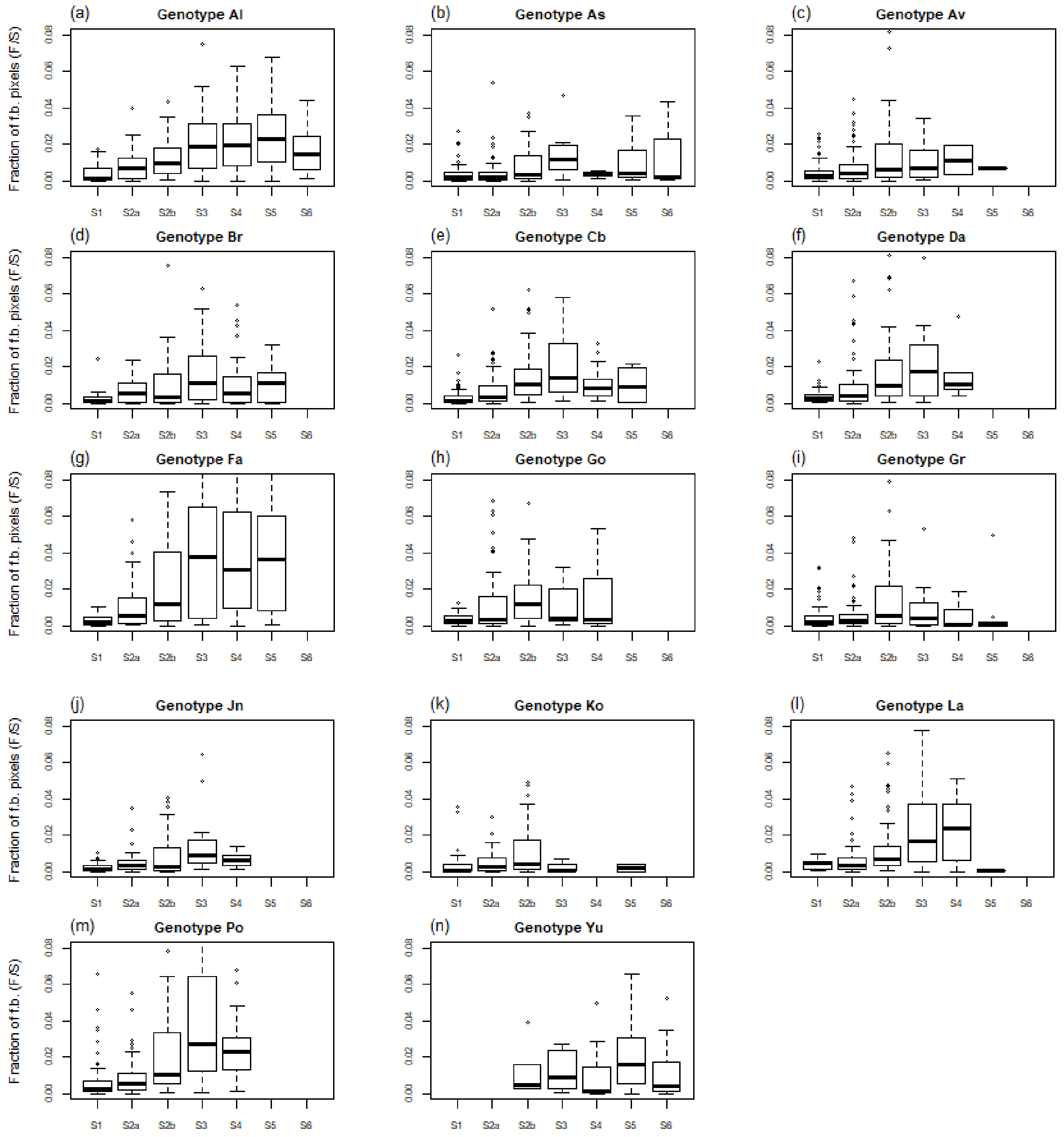

**Fig. S4.2.** Fraction of fruiting bodies pixels depending on the severity before incubation for the 15 genotypes in the Data1516 dataset.

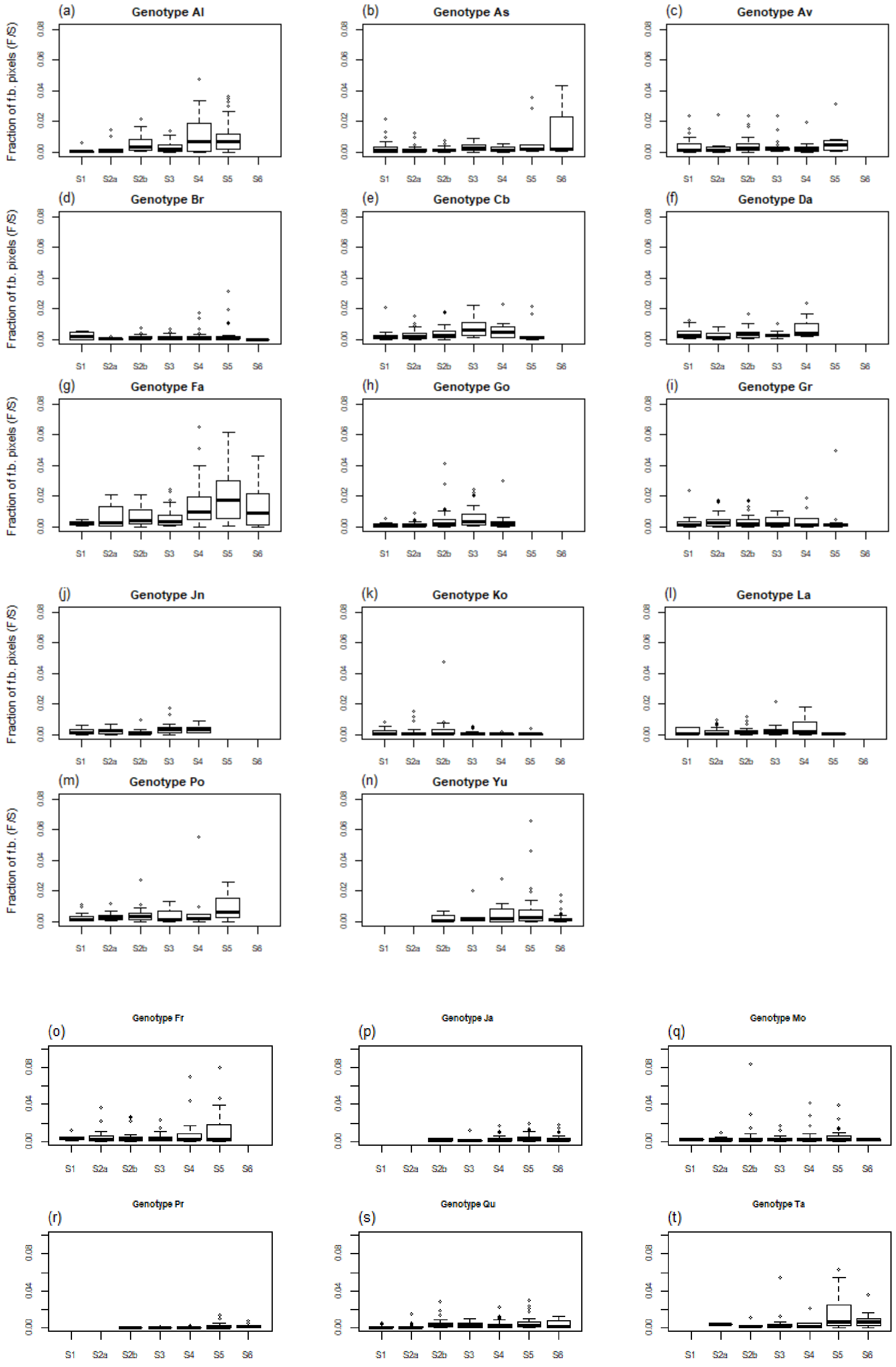

### S-5 Simulations

**Table S5.1** Parameters estimates of the proportional odds model with genotype effect used to predict event probabilities in the multinomial draw of simulations

| Coefficient | Estimate |
|---|---|
| S1\|S2a | -2,80 |
| S2a\|S2b | -1,48 |
| S2b\|S3 | -0,26 |
| S3\|S4 | 0,58 |
| S4\|S5 | 1,53 |
| S5\|S6 | 3,45 |
| Year1314 | 1,01 |
| Year1415 | 0,42 |
| Year1516 | 1,04 |
| NitrogenNl | -0,33 |
| GenotypeAs | -3,47 |
| GenotypeAv | -2,65 |
| GenotypeBr | -0,85 |
| GenotypeCb | -2,27 |
| GenotypeDa | -2,36 |
| GenotypeFa | -1,01 |
| GenotypeFr | -1,68 |
| GenotypeGo | -1,98 |
| GenotypeGr | -3,09 |
| GenotypeJa | 1,97 |
| GenotypeJn | -2,40 |
| GenotypeKo | -2,57 |
| GenotypeLa | -2,01 |
| GenotypeMo | -0,83 |
| GenotypeNh | -0,97 |
| GenotypePo | -2,50 |
| GenotypePr | 1,26 |
| GenotypeQu | -0,14 |
| GenotypeTa | 0,43 |
| GenotypeYu | 2,44 |
| NitrogenNl:GenotypeAs | 0,62 |
| NitrogenNl:GenotypeAv | 0,39 |
| NitrogenNl:GenotypeBr | 0,49 |
| NitrogenNl:GenotypeCb | 1,10 |
| NitrogenNl:GenotypeDa | 0,59 |
| NitrogenNl:GenotypeFa | 1,12 |
| NitrogenNl:GenotypeFr | 1,35 |
| NitrogenNl:GenotypeGo | 0,40 |
| NitrogenNl:GenotypeGr | 1,01 |
| NitrogenNl:GenotypeJa | 0,33 |
| NitrogenNl:GenotypeJn | -0,03 |
| NitrogenNl:GenotypeKo | 0,83 |
| NitrogenNl:GenotypeLa | 0,88 |
| NitrogenNl:GenotypeMo | 1,75 |
| NitrogenNl:GenotypeNh | 0,04 |
| NitrogenNl:GenotypePo | 1,10 |
| NitrogenNl:GenotypePr | 0,61 |
| NitrogenNl:GenotypeQu | -0,21 |
| NitrogenNl:GenotypeTa | 0,48 |
| NitrogenNl:GenotypeYu | 0,15 |

**Table S5.2** Parameters estimates of the proportional odds model without genotype effect used to predict event probabilities in the multinomial draw of simulations

| Coefficient | Estimate |
|---|---:|
| S1\|S2a | -1,05 |
| S2a\|S2b | 0,04 |
| S2b\|S3 | 0,93 |
| S3\|S4 | 1,49 |
| S4\|S5 | 2,09 |
| S5\|S6 | 3,43 |
| Year1314 | 0,81 |
| Year1415 | 0,38 |
| Year1516 | 0,88 |
| NitrogenNl | 0,24 |

**Table S5.3** Parameters estimates of GLM model with genotype effect used to predict the probability in the Binomial draw of simulations

| Coefficient | Estimate |
|---|---|
| (Intercept) | -5,12 |
| Year1314 | 0,49 |
| Year1415 | 0,16 |
| Year1516 | -1,11 |
| GenotypeAs | -0,52 |
| GenotypeAv | -0,33 |
| GenotypeBr | -0,62 |
| GenotypeCb | -0,23 |
| GenotypeDa | -0,04 |
| GenotypeFa | 0,41 |
| GenotypeFr | 0,55 |
| GenotypeGo | 0,10 |
| GenotypeGr | -0,37 |
| GenotypeJa | -0,95 |
| GenotypeJn | -0,50 |
| GenotypeKo | -0,56 |
| GenotypeLa | -0,24 |
| GenotypeMo | -1,16 |
| GenotypeNh | 0,04 |
| GenotypePo | 0,16 |
| GenotypePr | -1,73 |
| GenotypeQu | -0,71 |
| GenotypeTa | -0,04 |
| GenotypeYu | -0,48 |
| SeverityS2a | 0,50 |
| SeverityS2b | 1,04 |
| SeverityS3 | 1,30 |
| SeverityS4 | 1,29 |
| SeverityS5 | 1,52 |
| SeverityS6 | 1,23 |
| NitrogenNl | 0,71 |
| GenotypeAs:NitrogenNl | -0,24 |
| GenotypeAv:NitrogenNl | 0,03 |
| GenotypeBr:NitrogenNl | -0,59 |
| GenotypeCb:NitrogenNl | -0,11 |
| GenotypeDa:NitrogenNl | -0,47 |
| GenotypeFa:NitrogenNl | 0,71 |
| GenotypeFr:NitrogenNl | -1,02 |
| GenotypeGo:NitrogenNl | -0,69 |
| GenotypeGr:NitrogenNl | -0,09 |
| GenotypeJa:NitrogenNl | 0,03 |
| GenotypeJn:NitrogenNl | -0,43 |
| GenotypeKo:NitrogenNl | -1,41 |
| GenotypeLa:NitrogenNl | -0,49 |
| GenotypeMo:NitrogenNl | 1,48 |
| GenotypeNh:NitrogenNl | -0,76 |
| GenotypePo:NitrogenNl | -0,14 |
| GenotypePr:NitrogenNl | 0,24 |
| GenotypeQu:NitrogenNl | 0,25 |
| GenotypeTa:NitrogenNl | 0,77 |
| GenotypeYu:NitrogenNl | -0,44 |
| SeverityS2a:NitrogenNl | -0,51 |
| SeverityS2b:NitrogenNl | -0,82 |
| SeverityS3:NitrogenNl | -0,99 |
| SeverityS4:NitrogenNl | -0,69 |
| SeverityS5:NitrogenNl | -0,89 |
| SeverityS6:NitrogenNl | -1,21 |

**Table S5.4** Parameters estimates of GLM model without genotype effect used to predict the probability in the Binomial draw of simulations

| Coefficient | Estimate |
|---|---|
| (Intercept) | -5,32 |
| Year1314 | 0,49 |
| Year1415 | 0,25 |
| Year1516 | -1,22 |
| SeverityS2a | 0,54 |
| SeverityS2b | 1,08 |
| SeverityS3 | 1,41 |
| SeverityS4 | 1,45 |
| SeverityS5 | 1,60 |
| SeverityS6 | 1,12 |
| NitrogenNl | 0,54 |
| SeverityS2a:NitrogenNl | -0,52 |
| SeverityS2b:NitrogenNl | -0,85 |
| SeverityS3:NitrogenNl | -0,99 |
| SeverityS4:NitrogenNl | -0,65 |
| SeverityS5:NitrogenNl | -0,64 |
| SeverityS6:NitrogenNl | -0,61 |

**Fig. S5.1** Boxplot of simulated numbers of fruiting bodies pixels depending on the Year (2013 to 2016), on the Nitrogen level (Nh and Nl) and on the Genotype (21 varieties). The effect of Genotype was taken into account both for the distribution of stems in canker severity classes and on the prediction of fruiting bodies pixels, adjusting the model on observed data. Red bars are values without the Genotype effect. Data are means of 10 simulated field plots with 100 plants of 700 000 pixels.

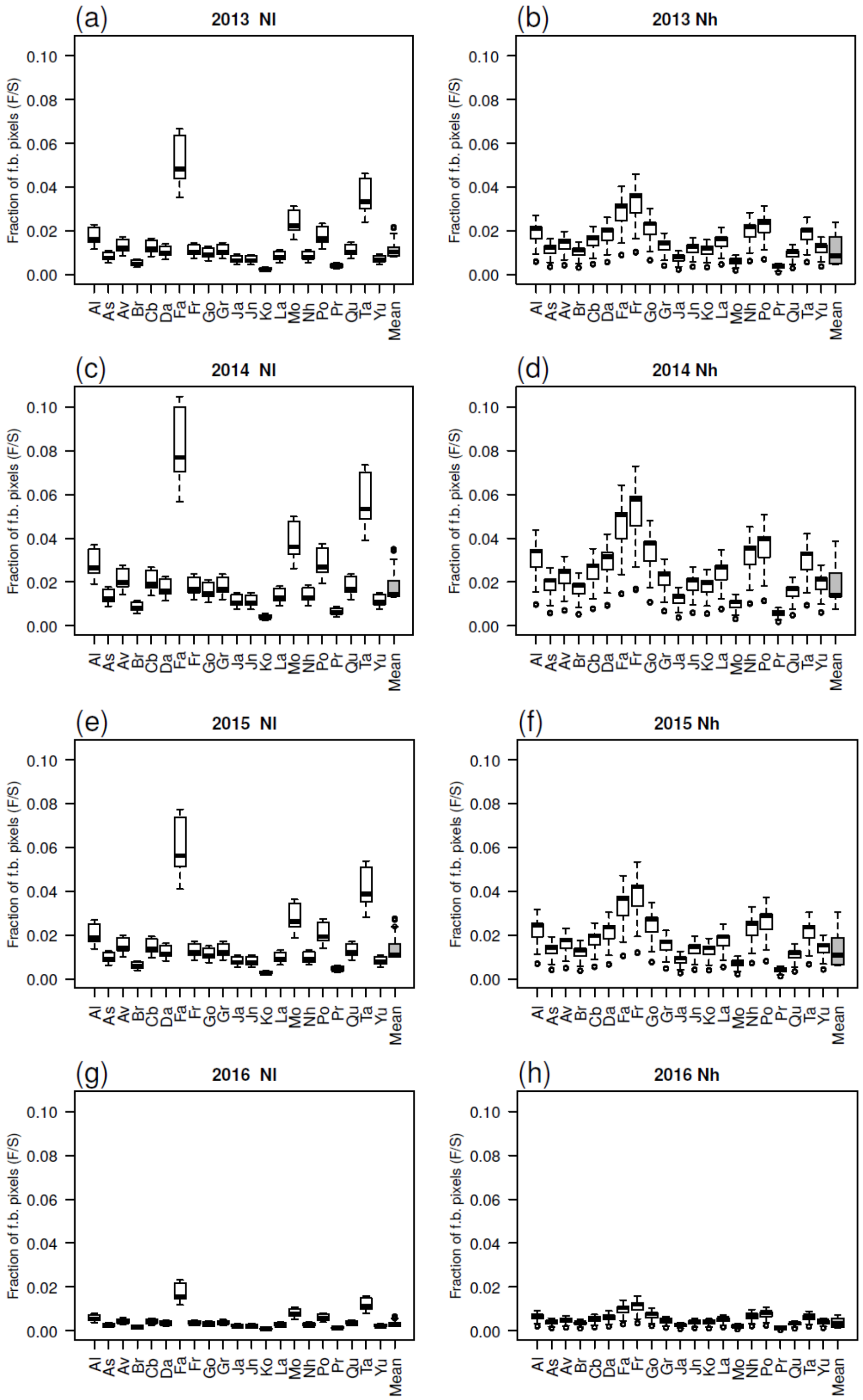
